# Drosophila STING protein has a role in lipid metabolism

**DOI:** 10.1101/2021.02.04.429825

**Authors:** Katarina Akhmetova, Maxim Balasov, Igor Chesnokov

## Abstract

Stimulator of interferon genes (STING) plays an important role in innate immunity by controlling type I interferon response against invaded pathogens. In this work we describe a direct but previously unknown role of STING in lipid metabolism in *Drosophila*. Flies with *STING* deletion are sensitive to starvation and oxidative stress, have reduced lipid storage and downregulated expression of lipid metabolism genes. We found that *Drosophila* STING interacts with lipid synthesizing enzymes acetyl-CoA carboxylase (ACC) and fatty acid synthase (FAS). ACC and FAS also interact with each other, indicating that all three proteins may be components of a large multi-enzyme complex. The deletion of *Drosophila STING* leads to disturbed ACC localization and decreased FAS enzyme activity. Together, our results demonstrate a direct role of STING in lipid metabolism in *Drosophila*.

## INTRODUCTION

STimulator of INterferon Genes (STING) is an endoplasmic reticulum-associated transmembrane protein that plays an important role in innate immune response by controlling the transcription of many host defense genes (Ishikawa and Barber 2008) (Ishikawa *et al.* 2009) (Sun *et al.* 2009) (Tanaka and Chen 2012) (Zhong *et al.* 2008). The presence of foreign DNA in the cytoplasm signals a danger for the cell. This DNA is recognized by specialized enzyme, the cyclic GMP-AMP synthase (cGAS) which generates cyclic dinucleotide (CDN) signaling molecules (Diner *et al.* 2013) (Li *et al.* 2013) (Gao *et al.* 2013) (Sun *et al.* 2013). CDNs bind to STING activating it (Wu *et al.* 2013) (Burdette *et al.* 2011) and the following signaling cascade results in NF-κB- and IRF3-dependent expression of immune response molecules such as type I interferons (IFNs) and pro-inflammatory cytokines (Sun *et al.* 2009) (Ishikawa *et al.* 2009) (Liu *et al.* 2015b). Bacteria that invade the cell are also known to produce CDNs that directly activate STING pathway (Sauer *et al.* 2011) (Woodward *et al.* 2010) (Danilchanka and Mekalanos 2013). Additionally, DNA which has leaked from damaged nuclei or mitochondria can also activate STING signaling and inflammatory response, which, if excessive or unchecked, might lead to the development of autoimmune diseases such as systemic lupus erythematosus or rheumatoid arthritis (Ahn *et al.* 2012) (Kawane *et al.* 2006) (Jeremiah *et al.* 2014) (Wang *et al.* 2015).

STING homologs are present in almost all animal phyla (Wu *et al.* 2014) (Margolis *et al.* 2017) (Kranzusch *et al.* 2015). This protein has been extensively studied in mammalian immune response, however, the role of STING in innate immunity of insects have been just recently identified (Hua *et al.* 2018) (Goto *et al.* 2018) (Liu *et al.* 2018) (Martin *et al.* 2018). Fruit fly *D.melanogaster STING* homolog is encoded by the *CG1667* gene, which we hereafter refer to as *dSTING.* dSTING displays anti-viral and anti-bacterial effects that however are not all encompassing. Particularly, it has been shown that dSTING-deficient flies are more susceptible to *Listeria* infection due to decreased expression of antimicrobial peptides (AMPs) - small positively charged proteins that possess antimicrobial properties against a variety of microorganisms (Martin *et al.* 2018). However, no effect has been observed during *E.coli* or *M.luteus* infections (Goto *et al.* 2018). dSTING has been shown to attenuate Zika virus infection in fly brains (Liu *et al.* 2018) and participate in the control of infection by two picorna-like viruses (DCV and CrPV) but not two other RNA viruses FHV and SINV or dsDNA virus IIV6 (Goto *et al.* 2018) (Martin *et al.* 2018). All these effects are linked to the activation of NF-κB transcription factor Relish (Kleino and Silverman 2014).

Immune system is tightly linked with metabolic regulation in all animals, and proper re-distribution of energy is crucial during immune challenges (Odegaard and Chawla 2013) (Alwarawrah *et al.* 2018) (Lee and Lee 2018). In both flies and humans, excessive immune response can lead to dysregulation of metabolic stores. Conversely, the loss of metabolic homeostasis can result in weakening of the immune system. The mechanistic links between these two important systems are integrated in *Drosophila* fat body (Arrese and Soulages 2010) (Buchon *et al.* 2014). Similarly to mammalian liver and adipose tissue, insect fat body stores excess nutrients and mobilizes them during metabolic shifts. Fat body also serves as a major immune organ by producing AMPs during infection. There is evidence that fat body is able to switch its transcriptional status from “anabolic” to “immune” program (Clark *et al.* 2013). The main fat body components are lipids, with triacylglycerols (TAGs) constituting approximately 90% of the stored lipids (Canavoso *et al.* 2001). Even though most of the TAGs stored in fat body comes from the dietary fat originating from the gut during feeding, *de novo* lipid synthesis in the fat body also significantly contributes to the pool of storage lipids (Heier and KÜhnlein 2018) (Wicker-Thomas *et al.* 2015) (Parvy *et al.* 2012) (Garrido *et al.* 2015).

Maintaining lipid homeostasis is crucial for all organisms. Dysregulation of lipid metabolism leads to a variety of metabolic disorders such as obesity, insulin resistance and diabetes. Despite the difference in physiology, most of the enzymes involved in metabolism, including lipid metabolism, are evolutionarily and functionally conserved between *Drosophila* and mammals (Lehmann 2018) (Toprak *et al.* 2020). Major signaling pathways involved in metabolic control, such as insulin system, TOR, steroid hormones, FOXO and many others, are present in fruit flies (Brogiolo *et al.* 2001) (Oldham *et al.* 2000) (JÜnger *et al.* 2003) (King-Jones and Thummel 2005). Therefore, it is not surprising that *Drosophila* has become a popular model system for studying metabolism and metabolic diseases (Teleman 2009) (Owusu-Ansah and Perrimon 2014) (Musselman and Kuhnlein 2018) (Baker and Thummel 2007) (Liu and Huang 2013) (Graham and Pick 2017) (Diop and Bodmer 2015). With the availability of powerful genetic tools, *Drosophila* has all the advantages to identify new players and fill in the gaps in our understanding of the intricacies of metabolic networks.

In this work we describe a novel function of dSTING in lipid metabolism. We report that flies with a deletion of *dSTING* are sensitive to starvation and oxidative stress. Detailed analysis reveals that *dSTING* deletion results in a significant decrease in the main storage metabolites, such as TAG, trehalose and glycogen. We identified two fatty-acid biosynthesis enzymes - Acetyl-CoA carboxylase (ACC) and fatty acid synthase (FAS) - as interacting partners for dSTING. Moreover, we also found that FAS and ACC interacted with each other, indicating that all three proteins might be components of a large complex. Importantly, *dSTING* deletion leads to decreased FAS activity and defects in ACC cellular localization suggesting a direct role of dSTING in lipid metabolism of fruit flies.

## RESULTS

### *Drosophila STING* mutants are sensitive to starvation and oxidative stress

Previously, we described a large genomic deletion that included *orc6* gene and the neighboring *CG1667 (dSTING)* gene which at that time was not characterized (Balasov *et al.* 2009). To create *dSTING* mutation we used the method of *P-*element imprecise excision. A *P*-element-based transposon *P{EPgy2} Sting*^*EY0649*^, located 353 base pairs upstream of the *dSTING* start codon, was excised by Δ2-3 transposase. *dSTINGΔ* allele contained deletion of 589 base pairs including start codon, first exon and part of the second exon (**Figure 1A**). Homozygotes *dSTINGΔ* mutant flies are viable with no obvious observable phenotype.

**Figure 1.**
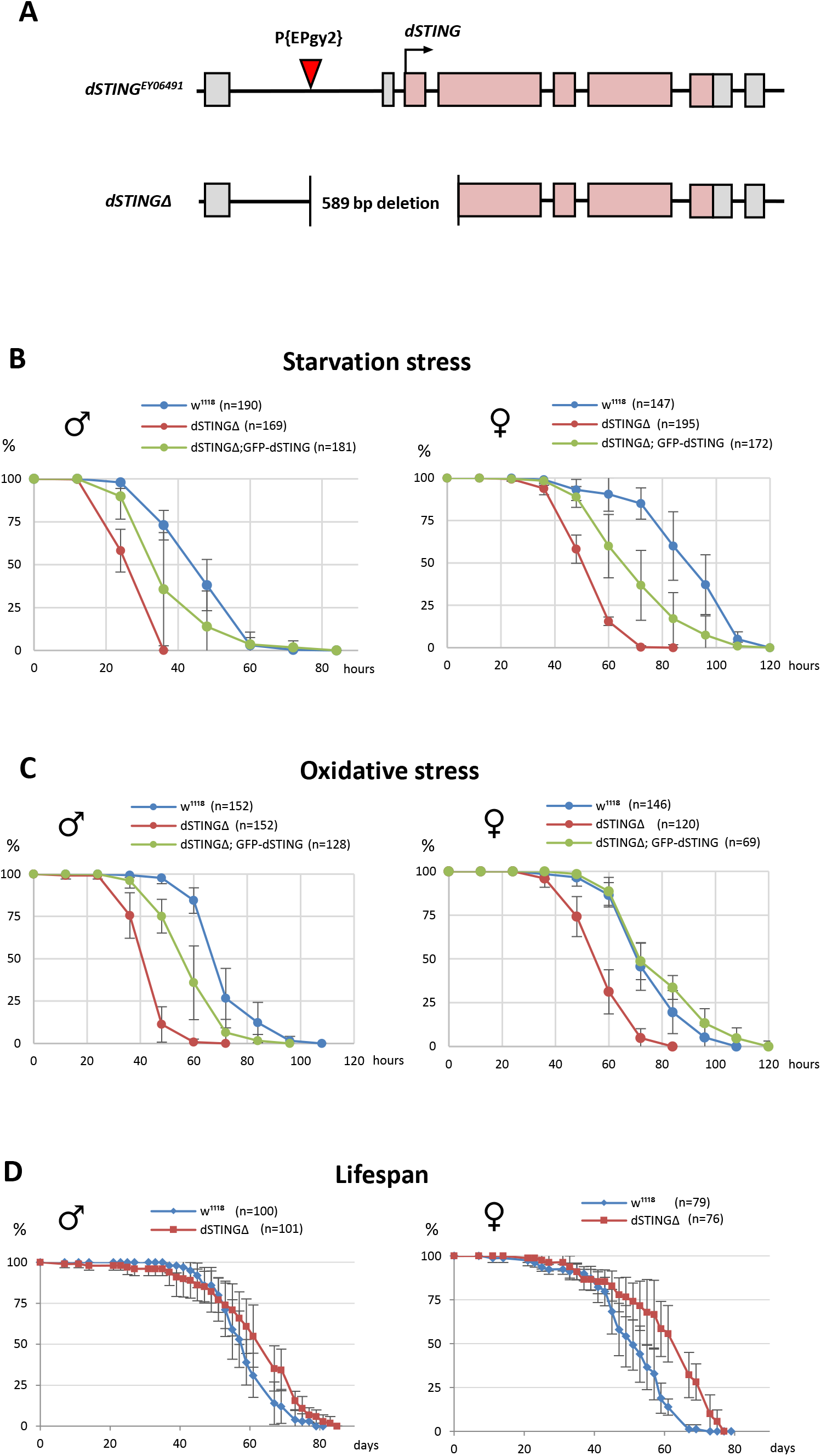
*Drosophila STING* mutants are susceptible to starvation and oxidative stress but have normal life span. **(A)** Generation of *Drosophila STING* deletion mutant. *dSTING* deletion mutant was generated by imprecise excision of P-element *P{EPgy2}STING*^*EY06491*^. *dSTINGΔ* allele contains a deletion of 589 base pairs including start codon, first exon and part of the second exon of *dSTING*. Exons are shown as pink-colored rectangles. The position of the P-element insertion is indicated by the red triangle. **(B)** Starvation stress resistance of males and females. Flies were kept on PBS only and percentages of survived flies were counted every 12 hours. **(C)** Oxidative stress resistance of males and females. Flies were kept on food supplemented with 5% hydrogen peroxide and percentages of survived flies were counted every 12 hours. **(D)** Lifespan of males and females. Flies were kept on regular food and percentages of survived flies were counted. **(B-D)** Percentages of survived flies at each time point are shown. The number of flies analyzed is shown in chart legend for each genotype.

The role of dSTING in anti-viral and anti-bacterial defense in *Drosophila* has been established recently (Martin *et al.* 2018) (Goto *et al.* 2018) (Liu *et al.* 2018). Since the changes in immune response are often accompanied by a dysregulation of metabolic homeostasis and vice versa (Zmora *et al.* 2017) (Odegaard and Chawla 2013) (Alwarawrah *et al.* 2018), we analyzed *dSTINGΔ* mutant flies for the defects in metabolism. A response to the metabolic stress is a good indicator of defects in metabolism, therefore we subjected flies to starvation stress and oxidative stress. We found that *dSTINGΔ* mutant flies were sensitive to both starvation and oxidative stress as compared to the control flies (**Figure 1B,C**). *dSTINGΔ* mutant larvae were also more susceptible to both types of stress (**Supplementary Figure 1A,B**). To confirm that the observed phenotypes are not due to off-target effects, we designed fly strain containing GFP-tagged wild type *dSTING* (under the native *dSTING* promoter) on *dSTINGΔ* deletion background. The expression of *GFP-dSTING* partially or entirely rescued the sensitivity of *dSTINGΔ* deletion flies to both starvation and oxidative stress (**Figure 1B,C**), suggesting that observed phenotypes are caused by dSTING deficiency. Interestingly, the deletion of *dSTING* had no effect on the total lifespan of fed flies in both males and females. Moreover, the age-related mortality was slightly reduced, especially for females (**Figure 1D**).

It is possible that increased sensitivity to starvation and oxidative stress that we observed in *dSTINGΔ* flies is caused by decreased defense against commensal or pathogenic bacteria in the absence of *dSTING*. To test this hypothesis, we generated axenic, or germ-free, flies. We found that under axenic condition *dSTINGΔ* mutants exhibited the same response to starvation and oxidative stress as *dSTINGΔ* non-axenic flies (**Supplementary Figure 2**), suggesting that diminished immune response against bacteria is not likely to be responsible for the observed phenotypes.

Collectively, our data suggest that deletion of *Drosophila STING* results in increased susceptibility of flies to starvation and to oxidative stress.

### *Drosophila STING* mutants have decreased storage metabolites

The ability of an organism to store nutrients when they are abundant is crucial for its survival during periods of food shortage. Triacylglycerols (TAGs), glycogen and trehalose are major metabolites for carbon storage in *Drosophila*. Dietary glucose absorbed from the gut is quickly converted to trehalose, which is a main hemolymph sugar in insects. Glycogen is another form of carbohydrate storage that accumulates in the fat body and muscles. Finally, most energy reserves in insects are in the form of lipids, particularly TAGs that are stored in lipid droplets of the fat body (Canavoso *et al.* 2001).

We measured storage metabolites levels along with glucose level in fed or 24 hour starved adult males. Under fed conditions, TAG level was decreased two-fold in *dSTINGΔ* mutants as compared to control flies. Under starved condition, TAG level dropped dramatically to about 1/8 of the wild type level (**Figure 2A**). Glycogen and trehalose levels were also significantly decreased in *dSTINGΔ* mutants in both fed and starved flies (**Figure 2B,C**). Interestingly, glucose level was increased under fed condition (**Figure 2D**), suggesting that *dSTINGΔ* mutant flies have either decreased incorporation of ingested glucose into trehalose or glycogen, or increased breakdown of these storage molecules. Nevertheless, when flies were starved for 24 hours, glucose level in *dSTINGΔ* mutants dropped and was two-fold lower than in control flies (**Figure 2D, starved**). In fed flies, the expression of GFP-tagged *dSTING* partially rescued the mutant phenotypes for all measured metabolites. However, under starved condition, rescue was observed only for TAG level (**Figure 2**).

**Figure 2.**
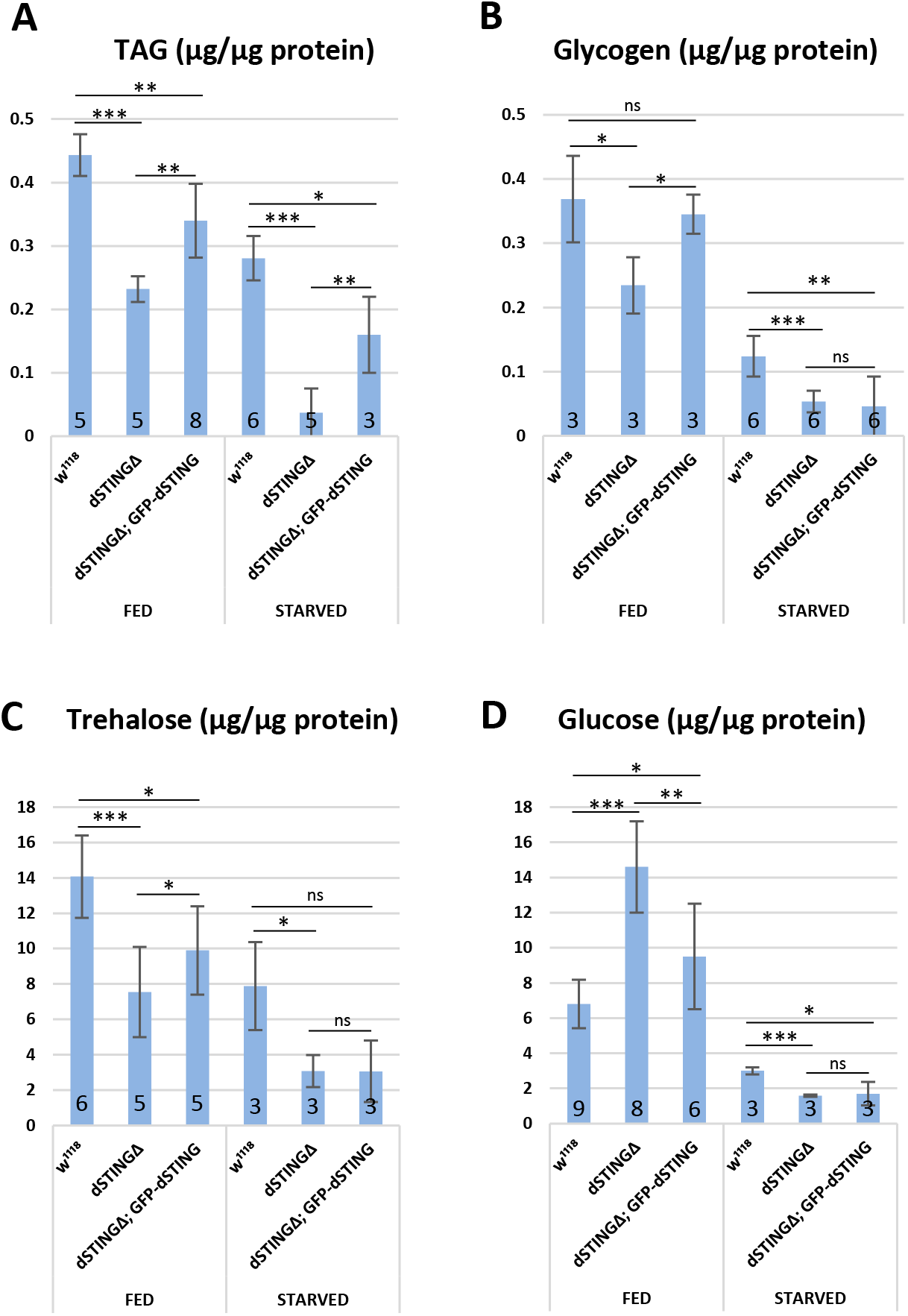
Storage metabolites are significantly decreased in *Drosophila STING* mutants. Metabolites were measured in fed or 24 hours starved males. **(A), (B)** TAG and glycogen levels were measured in the total body. **(C), (D**) Trehalose and glucose levels were measured in the hemolymph. Levels of metabolites are shown per μg of total protein. The number of experiments for each genotype is indicated. Student’s t-test, *p<0.05, **p<0.01, ***p<0.001, ns indicates statistically non significant.

Two RNAi screens for obesity and anti-obesity genes in *Drosophila* did not reveal any significant changes in TAG level in dSTING-deficient flies (Pospisilik *et al.* 2010) (Baumbach *et al.* 2014). The potential discrepancy with our data might be explained by the fact that in both mentioned studies RNAi was induced only 2-8 days after eclosion, whereas in our study dSTING was absent from the very beginning of the development.

One of the possible explanations for the decreased storage metabolites might be a decrease in food consumption. To test this possibility, we used capillary feeder (CAFE) assay (Ja *et al.* 2007) which showed that it was not the case, and *dSTINGΔ* mutant flies consumed food at the same rate as control flies (**Supplementary Figure 3A**).

Also, a compromised gut barrier function could potentially lead to decreased nutrient absorption and susceptibility to starvation stress. To assess intestinal permeability *in vivo* we performed “smurf” assay (Rera *et al.* 2012). Flies were fed blue dye and checked for the presence of the dye outside of the digestive tract. “Smurf” assay did not reveal any loss of gut wall integrity in *dSTINGΔ* mutants (**Supplementary Figure 3B**).

Together, these data indicate that a deletion of *dSTING* results in a disruption of lipid/carbohydrate energy balance.

### Lipid metabolism is impaired in *Drosophila STING* mutants

Among measured metabolites, the effect of dSTING mutation on TAG level was the most pronounced. Moreover, the expression of *GFP-dSTING* on *dSTINGΔ* mutant background partially rescued TAG levels under both fed and starved conditions (**Figure 2**). Therefore, we decided to take a closer look at the lipid metabolism in the absence of dSTING. In insects, TAG lipids are stored mainly in fat body and midgut in the form of cytoplasmic lipid droplets. To visualize lipid stores, we stained fat bodies and midguts of adult flies with Nile Red dye that selectively labels lipids within the cells (**Figure 3**). Fat bodies of *dSTINGΔ* mutant flies contained significantly fewer lipids as compared to the control flies (**Figure 3A,A’**). The expression of *GFP-dSTING* rescued this phenotype. *dSTINGΔ* mutant larvae also had decreased lipid droplet content in fat body (**Supplementary Figure 1C**). Interestingly, lipid droplet content in midguts was not decreased in *dSTINGΔ* mutants (**Figure 3B,B’**) indicating that only fat body lipid storage was affected. In support of these data, fat body specific RNAi of *dSTING* resulted in starvation sensitivity and reduced TAG level in adult flies (**Supplementary Figure 4**), highlighting the role of dSTING in fat body functions.

**Figure 3.**
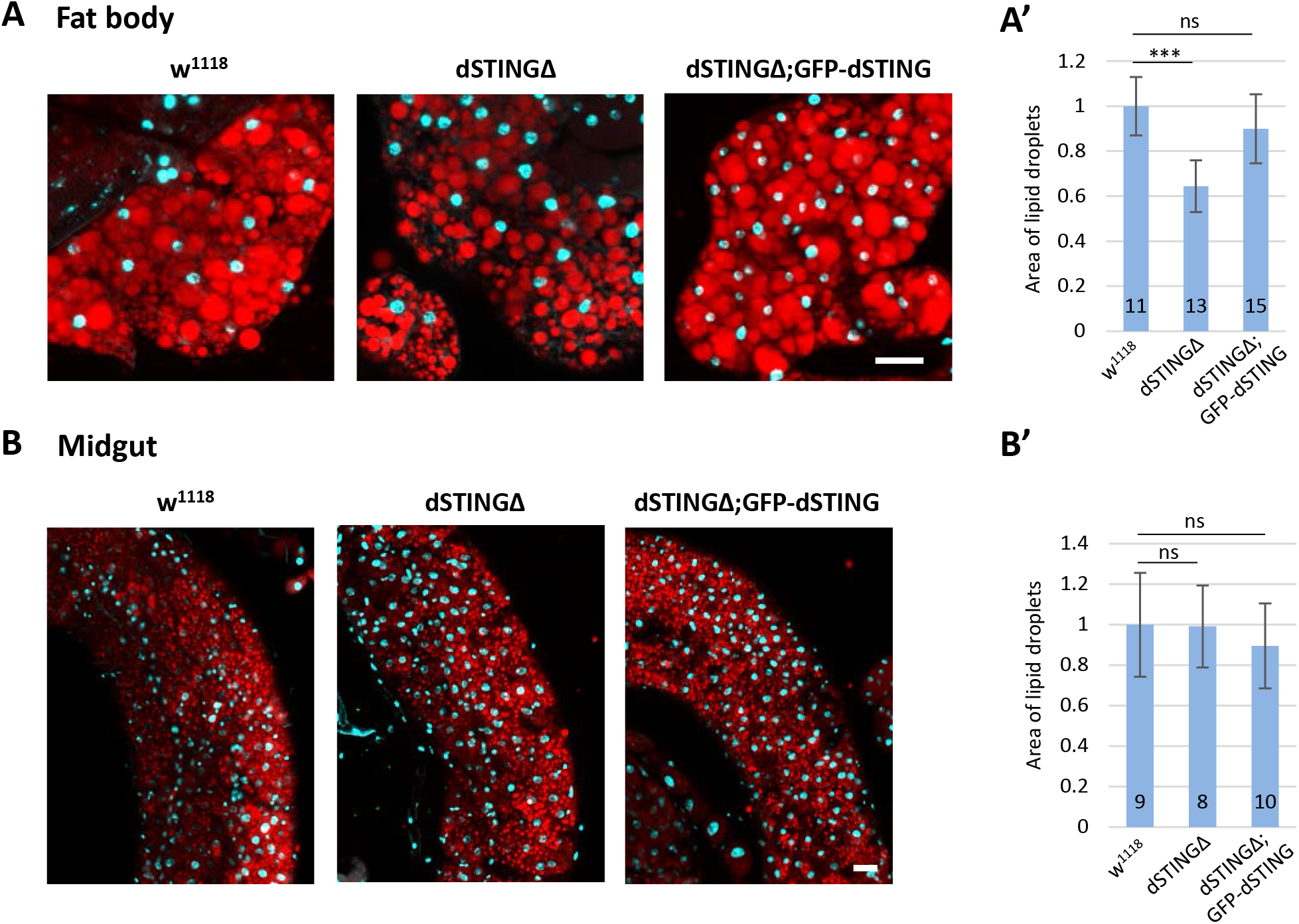
Drosophila *STING* mutants have decreased fat body lipid storage. Adult fat body **(A)** or midguts **(B)** were stained with Nile Red (red) that labels lipid droplets. Nuclei were stained with DAPI (blue). Scale bar 20μm. **(A’) (B’)** Quantification of lipid droplets area. Values were normalized to wild type (*w*^*1118*^). Number of samples analyzed is shown for each genotype. Student’s t-test, ***p<0.001, ns indicates statistically non significant.

Nutrient deprivation as well as other cellular stress conditions, such as damaged organelles and infection, induce autophagy response (Mizushima *et al.* 2002) (Deretic and Levine 2018). STING has been linked to autophagy on various levels (Moretti *et al.* 2017) (Bhatelia *et al.* 2017) (Liu *et al.* 2019). In *Drosophila*, activation of autophagy is of particular importance since it serves as the response to nutrient starvation, and mutants with defects in autophagy are hypersensitive to starvation (JuhÁsz *et al.* 2007). However, we did not observe a reduction of autophagy in fat bodies of *dSTINGΔ* mutants (**Supplementary Figure 5**). We cannot exclude a possibility that autophagy response to another stimuli or in different tissues is defective in the absence of dSTING. For example, dSTING was shown to attenuate Zika virus infection in fly brains through the induction of autophagy pathways (Liu *et al.* 2018).

To gain insight into gene expression changes in the absence of *dSTING*, we performed microarray analysis of *dSTINGΔ* mutant and control flies under fed and 24 hour starved conditions. Under fed conditions, microarray analysis revealed a significant change in 672 transcripts (more than 1.4 fold change), with 381 transcripts expressed at reduced levels and 291 transcripts at elevated levels. Under starved conditions, the expression of 1452 genes was altered in *dSTINGΔ* mutants, with 797 downregulated and 655 upregulated genes (**Supplemental Table 1**).

Principal components analysis (PCA) is a common method for the analysis of gene expression data, providing information on the overall structure of the analyzed dataset (Lever *et al.* 2017). PCA plot for our microarray data showed that the sample groups separated along the PC1 axis (which explained 29% of all variance in the experiment), with the greatest separation between the wild-type fed and mutant starved groups (**Figure 4A**). Interestingly, the *dSTINGΔ* fed and the wild-type starved groups clustered together along PC1 axis, indicating that *dSTING* knockout and wild type flies starvation induced similar changes in gene expression.

**Figure 4.**
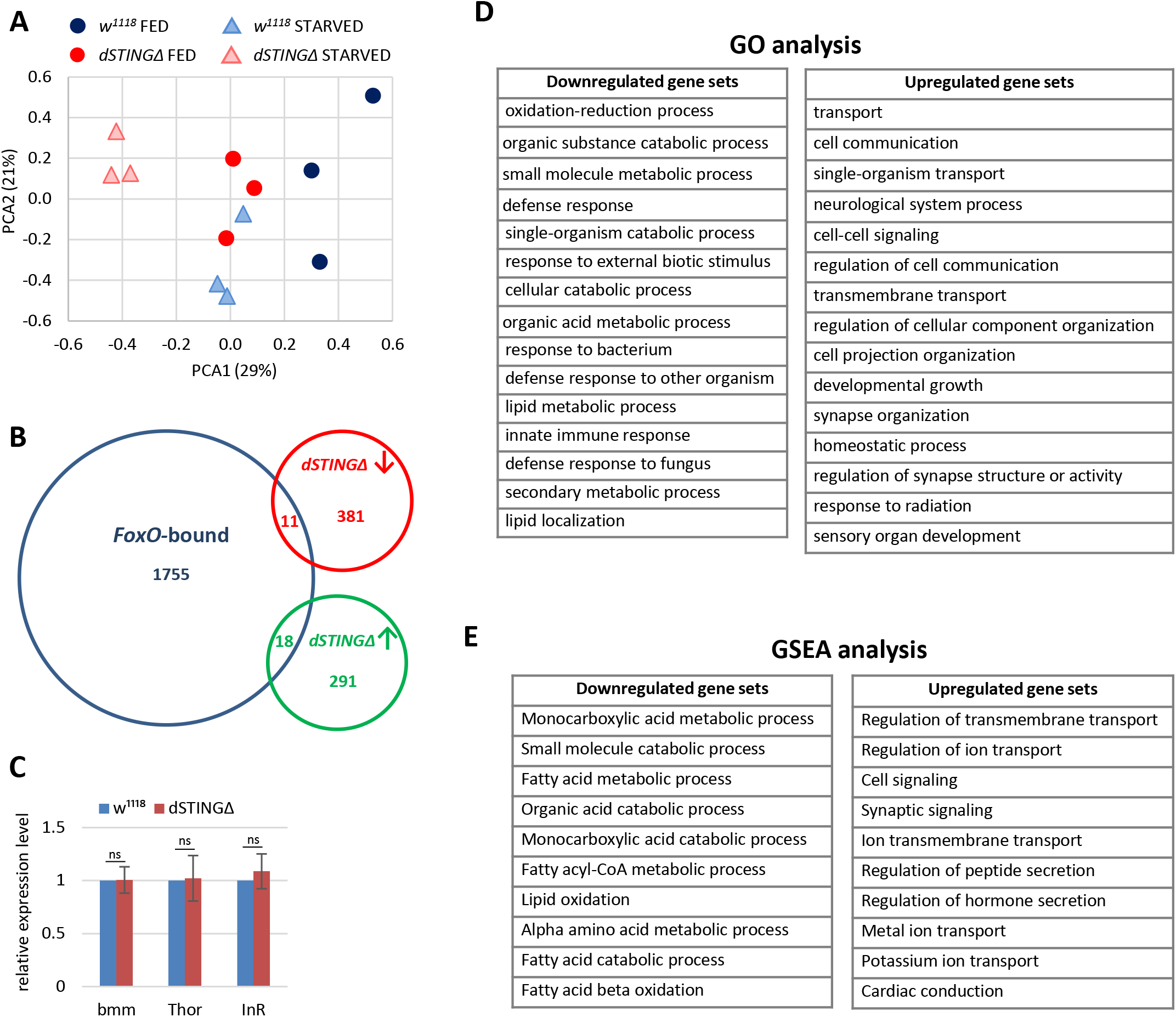
Lipid metabolism genes are downregulated in *Drosophila STING* mutants. Fed or 24 hours starved adult males (*dSTINGΔ* mutants or *w*^*1118*^ as a control) were subjected to microarray analysis. **(A)** Principal component analysis (PCA) of microarray data. PCA scores plot showing variances in gene expression profiles between groups is shown. Each sample is shown as a single point (n=3 per genotype). **(B)** Overlap between the genes bound by *Drosophila* FoxO {Alic, 2011 #116} {Teleman, 2008 #117} and genes downregulated or upregulated in *dSTINGΔ* mutants. **(C)** Transcription of FoxO targets measured by RT-qPCR. The bars indicate relative expression level. Values were normalized to wild type (*w*^*1118*^). Student’s t-test, ns indicates statistically non significant. **(D)** Gene ontology (GO) analysis of microarray data. *dSTINGΔ* mutants and control *w*^*1118*^ under fed conditions were compared. Downregulated and upregulated top scoring gene sets are shown. **(E)** Gene Set Enrichment Analysis (GSEA) of microarray data. *dSTINGΔ* mutants and control *w*^*1118*^ under fed conditions were compared. Downregulated and upregulated top scoring gene sets are shown.

It was shown previously that the increased activity of FoxO transcription factor can phenocopy starvation conditions in *Drosophila* (Molaei *et al.* 2019) (Kramer *et al.* 2003). FoxO is also known to induce the expression of various genes encoding proteins involved in lipid mobilization (Teleman *et al.* 2005) (Wang *et al.* 2011). Therefore, the decreased TAG phenotype and gene expression pattern resembling starvation in *dSTINGΔ* mutants could be explained by FoxO activation. However, we found that only a small fraction of genes controlled by FoxO was upregulated in *dSTINGΔ* mutants (**Figure 4B**) (Alic *et al.* 2011) (Teleman *et al.* 2008). Particularly, the expression of main FoxO targets, including lipase *brummer*, translational regulator *Thor* and insulin receptor *InR* was not affected in the absence of dSTING (**Figure 4C**). Consequently, we did not observe the increase in nuclear FoxO which is a characteristic of FoxO activation (**Supplementary Figure 6**).

In agreement with a report by Martin et al. (Martin *et al.* 2018), *dSTINGΔ* mutants are characterized by the downregulation of immune response genes, including AMPs (*Mtk*, *Drs*, *AttD*, *DptB*, *BomS1*), peptidoglycan recognition proteins (PGRPs, such as *PGRP-SD* and *PGRP-SA)*, and serpins - (*Spn53F*, *Spn42De*) (**Supplemental Table 1**). These results are expected since *STING* was initially discovered in fruit flies and silkworm as an immune response gene (Martin *et al.* 2018), (Hua *et al.* 2018), (Goto *et al.* 2018). To gain more insight into the biological processes that are altered in the absence of dSTING, we looked at the gene set enrichment in *dSTINGΔ* mutants under fed conditions. Based on the Gene Ontology (GO) analysis, metabolic processes and immune response genes were downregulated in *dSTINGΔ* mutants (**Figure 4D, downregulated gene sets**). Upregulated genes were enriched with GO classifications related to cell signaling (e.g. transport, cell communication and synapse organization) (**Figure 4D, upregulated gene sets**).

GO analysis requires a discrete list of genes (downregulated and upregulated in our case). Gene Set Enrichment Analysis (GSEA) on the other hand, uses all microarray datapoints, therefore it is expected to be more sensitive since it can identify gene sets comprising many members that are undergoing subtle changes in expression (Subramanian *et al.* 2005). We analyzed *dSTINGΔ* mutants versus control flies under fed conditions using GSEA approach and found that metabolism of lipids, particularly fatty acids, was among top scoring gene sets downregulated in *dSTINGΔ* mutants (**Figure 4E, downregulated gene sets**). As for upregulated gene sets, GSEA data were similar to GO analysis data (**Figure 4E, upregulated gene sets**).

Together, we found that *Drosophila STINGΔ* mutants have defects in lipid metabolism manifested in decreased lipid storage in fat body and decreased expression of lipid metabolism genes.

### *Drosophila STING* protein interacts with Acetyl-CoA Carboxylase and Fatty Acid Synthase

In mammals, STING is an adaptor molecule that activates downstream signaling through protein-protein interactions. To look for possible interaction partners for *Drosophila* STING that could explain its effect on lipid metabolism, we performed immunoprecipitation from fat bodies of larvae expressing *GFP-dSTING* using anti-GFP antibody. Immunoprecipitated material was separated by SDS-PAGE, and the most prominent bands were subjected to mass spectrometry analysis. Several proteins with a high score were identified, including Fatty acid synthases 1 and 2 (CG3523 and CG3524, respectively), Acetyl-CoA carboxylase (CG11198) and dSTING itself (CG1667) (**Supplementary Table 2**).

Acetyl-CoA carboxylase (ACC) and fatty acid synthase (FAS) are two important enzymes of the *de novo* lipid biosynthesis pathway (LÓpez-Lara and Soto 2019) (Wakil and Abu-Elheiga 2009). ACC catalyzes the formation of malonyl-CoA from acetyl-CoA, the first committed step of fatty acid synthesis. The next step is performed by FAS which uses malonyl-CoA and acetyl-CoA to synthesize palmitic fatty acid. Palmitate might undergo separate elongation and/or unsaturation by specialized enzymes to yield other fatty acid molecules. A series of reactions then add the fatty acids to a glycerol backbone to form triacylglycerol (TAG), the main energy storage molecule.

We confirmed the mass spec results by performing immunoprecipitation from abdomens of adult flies expressing *GFP-dSTING* in fat body. Both ACC and FAS co-immunoprecipitated with GFP-dSTING (**Figure 5A, B**). Interestingly, we found that ACC and FAS interacted with each other as showed by the reciprocal immunoprecipitation experiments (**Figure 5C, D**). We observed this interaction not only in control flies but also in *dSTINGΔ* mutants flies with or without the expression of *GFP-dSTING*.

**Figure 5.**
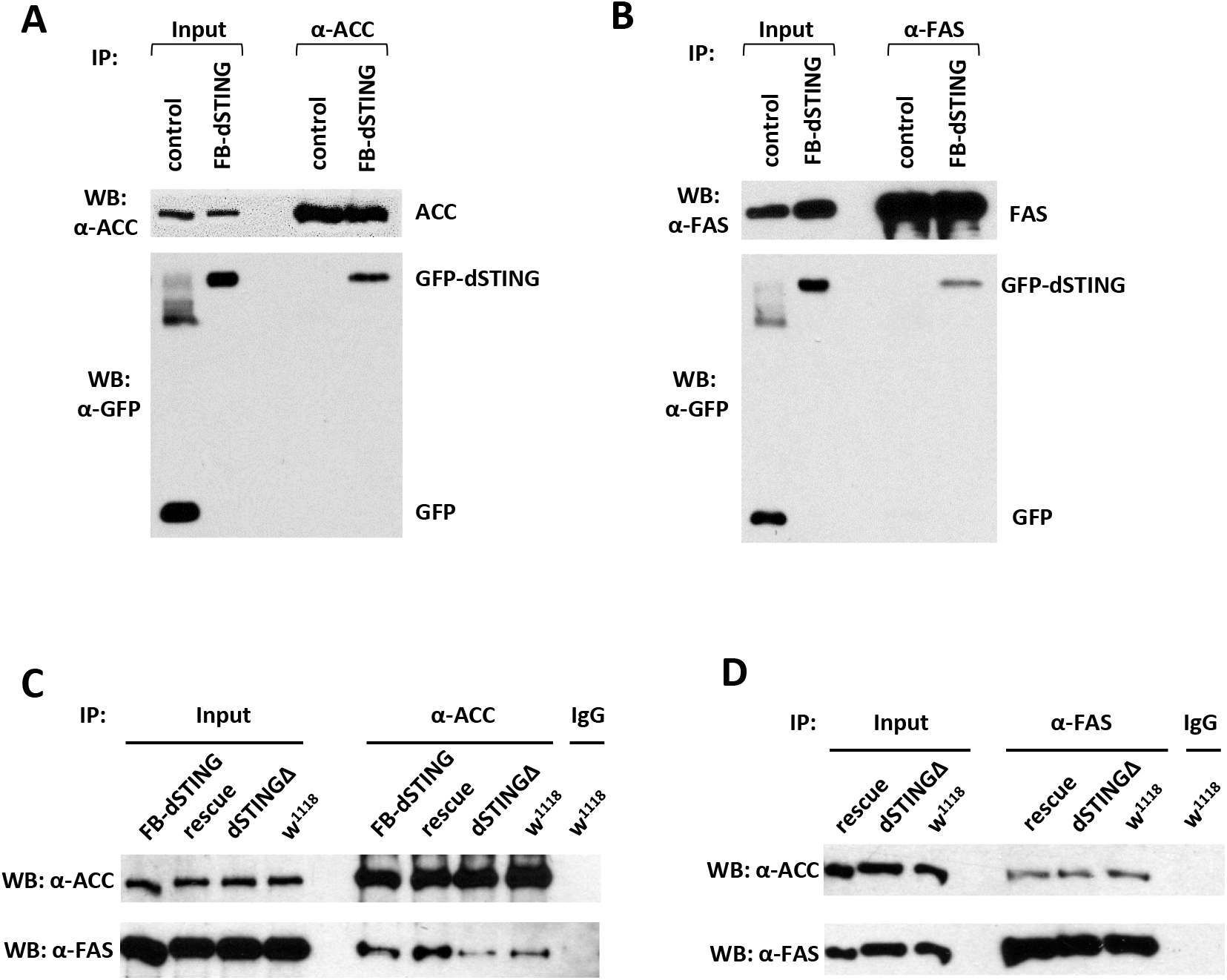
*Drosophila* STING protein interacts with acetyl-CoA carboxylase (ACC) and fatty acid synthase (FAS). **(A) (B)** dSTING interacts with ACC and FAS. ACC **(A)** or FAS **(B)** were immunoprecipitated from abdomens of adult flies using corresponding antibody. Genotypes are: control – *w*^*1118*^, FB-dSTING – *cg-GAL4/GFP-dSTING* (flies expressing GFP-dSTING in fat body). Recombinant GFP was added to the control reaction. **(C) (D)** ACC and FAS interact with each other. ACC **(C)** or FAS **(D)** were immunoprecipitated from abdomens of adult flies using corresponding antibody. Rescue – *dSTINGΔ;GFP-dSTING,* FB-dSTING – *cg-GAL4/GFP-dSTING*.

Together, our data indicate that dSTING, ACC and FAS interact with one another, suggesting that they might components of a multi-protein complex involved in fatty acid synthesis.

### Fatty acid synthase activity is decreased in *Drosophila STING* mutants

Since dSTING was found as part of a complex with ACC and FAS, we asked if *dSTING* deletion might result in changes in these enzyme activities. We measured ACC and FAS activity in adult flies. ACC protein level and activity were not significantly changed in *dSTINGΔ* mutants flies (**Figure 6A,A’**). However, FAS activity was almost two times lower in *dSTINGΔ* mutants as compared to the control flies (**Figure 6B**), but the protein level was unchanged (**Figure 6B’**). Importantly, the expression of GFP-tagged dSTING on *dSTINGΔ* mutant background restored FAS activity to control level (**Figure 6B**).

**Figure 6.**
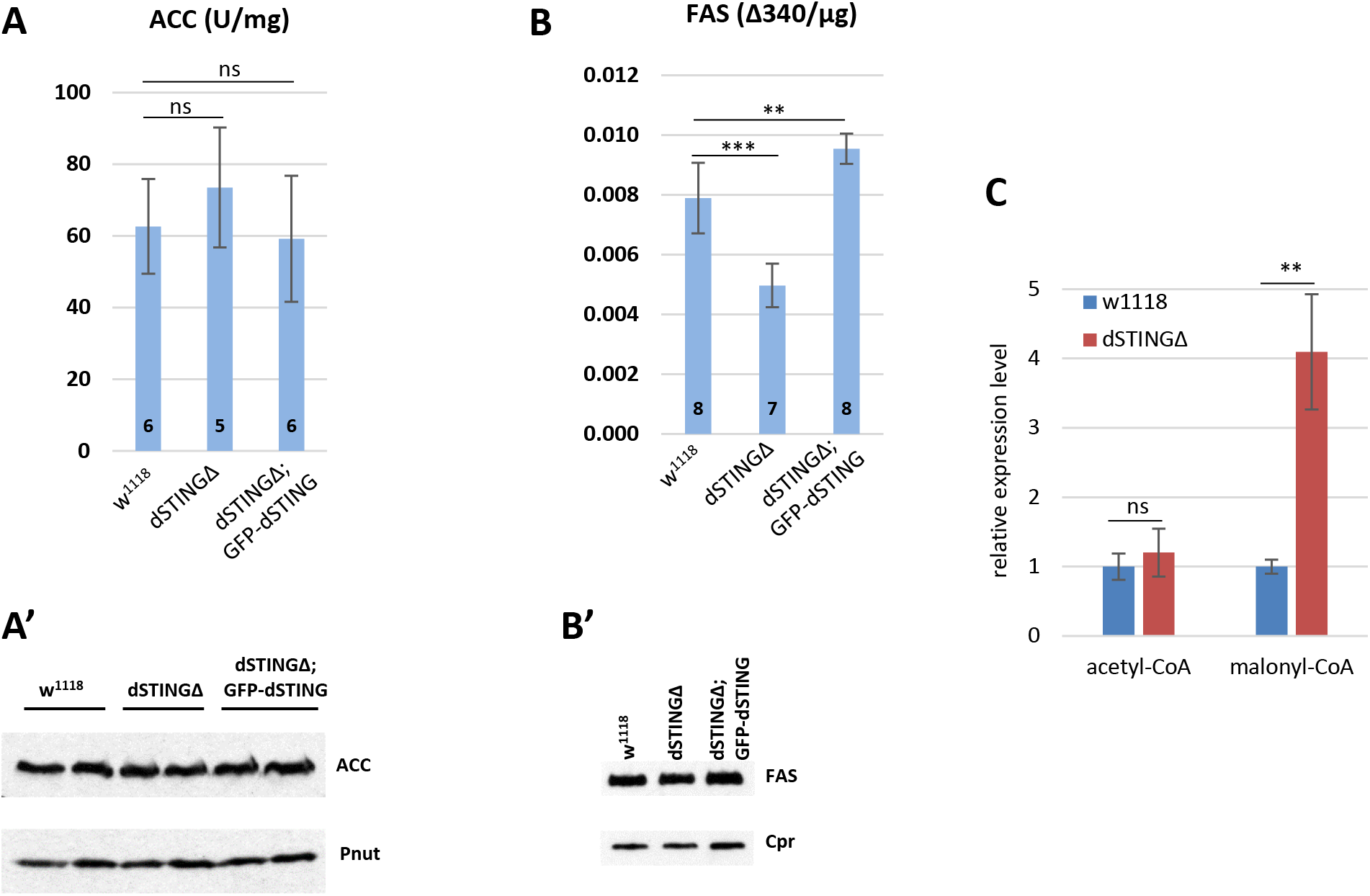
Fatty acid synthase activity is decreased in *Drosophila STING* mutants. **(A)(B)** Enzyme activity assays. ACC activity **(A)** or FAS activity **(B)** was measured in the total body of adult flies and normalized to protein level. The number of experiments for each genotype is indicated. Student’s t-test, **p<0.01, ***p<0.001, ns indicates statistically non significant. **(A’)** ACC protein level in total fly extract. Pnut was used as a loading control. **(B’)** FAS protein level in total fly extract. Cpr was used as a loading control. **(C)** Acetyl-CoA and malonyl-CoA levels in fly total body extracts. Values were normalized to wild type (*w*^*1118*^). Student’s t-test, ***p<0.01, ns indicates statistically non significant.

ACC enzyme carboxylates acetyl-CoA resulting in the formation of malonyl-CoA which then serves as a substrate for FAS in the synthesis of fatty acids. If ACC activity is unchanged and FAS activity is decreased we should observe the accumulation of malonyl-CoA. Polar metabolite profiling of *dSTINGΔ* flies compared to control flies showed that indeed, malonyl-CoA level was significantly increased whereas acetyl-CoA level remained unchanged in the mutants (**Figure 6C**, **Supplementary Figure 7**).

### ACC localization is perturbed in the fat body of *dSTINGΔ* mutants

In mammals, STING is a transmembrane protein that localizes to endoplasmic reticulum (ER). To check whether this is also the case in *Drosophila*, we performed membrane fractionation, which showed that GFP tagged dSTING co-sedimented exclusively with membrane fraction (**Supplementary Figure 8A**). We also expressed *GFP-dSTING* in *Drosophila* S2 tissue culture cells and found that it mostly co-localized with ER and to the lesser extent with Golgi, but not with cellular membrane (**Supplementary Figure 8B**), in agreement with previous observation (Goto *et al.* 2018).

Next, we used adult flies expressing GFP-tagged dSTING under native *dSTING* promoter to examine the localization of dSTING, ACC and FAS in the fat body, main lipid synthesizing organ in *Drosophila.* ER (as judged by ER marker Calnexin) extended throughout fat body cells, with most prominent staining at the cell periphery and in perinuclear region as was shown before (Jacquemyn *et al.* 2020) (**Figure 7A**). GFP-dSTING mainly co-localized with Calnexin at the cortex (**Figure 7A**). Little or no signal was observed at the perinuclear region of fat body cells. Both ACC and FAS partially co-localized with GFP-dSTING at the cell periphery region of ER (**Figure 7B,C**).

**Figure 7.**
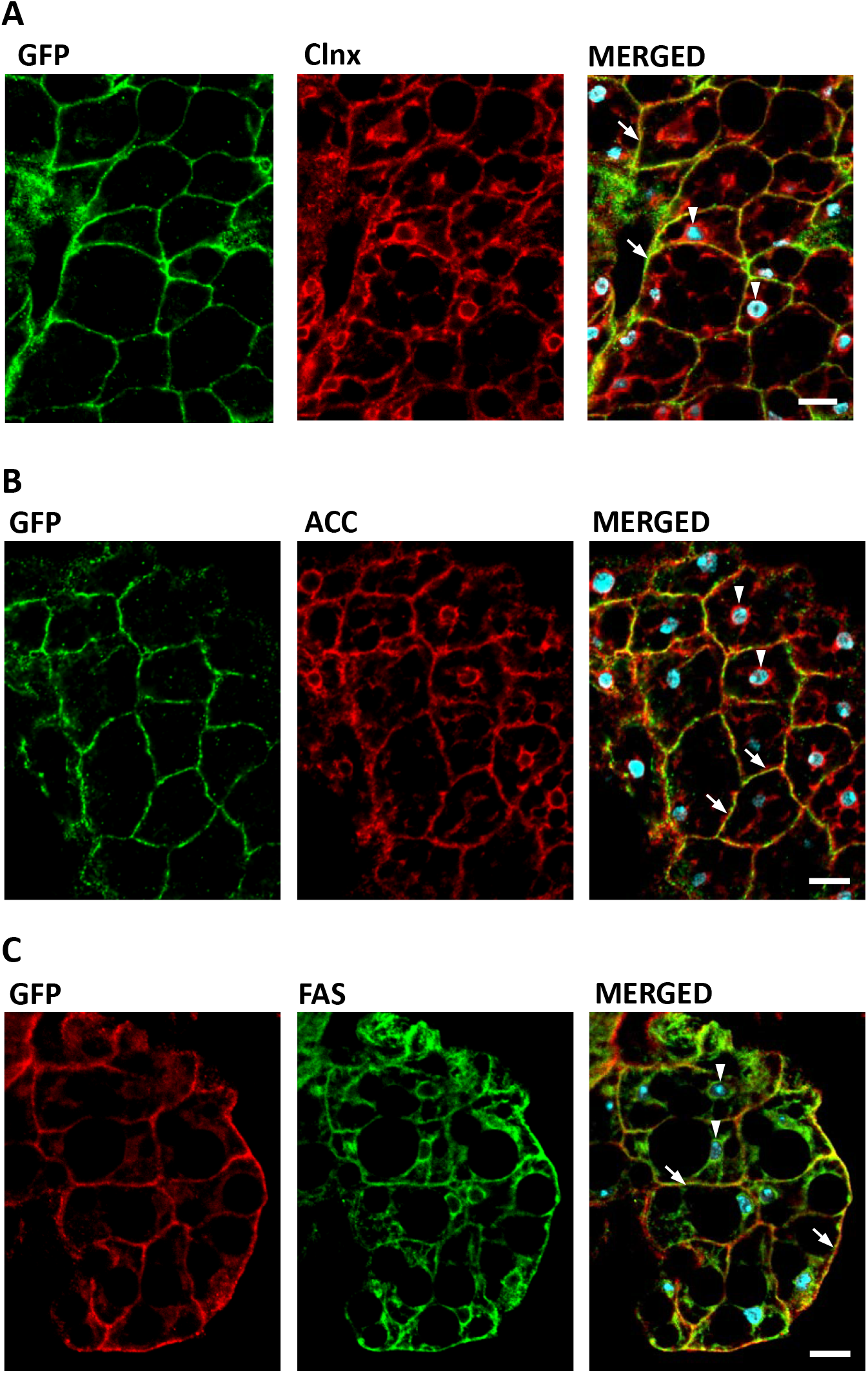
dSTING, ACC and FAS co-localize in *Drosophila* fat body cells. Fat body of adult flies expressing GFP-tagged dSTING (genotype *dSTINGΔ;GFP-dSTING*) were stained for: **(A)** GFP (green) and Calnexin (Clnx, red); **(B)** GFP (green) and ACC (red); **(C)** GFP (red) and FAS (green). Nuclei were stained with DAPI (blue). Arrows mark cortical region, arrowheads mark perinuclear region of fat body cells. Scale bar 10μm.

We asked whether dSTING mutation affected localization of ACC or FAS in fly fat body. While ACC extended throughout wild-type cells, in *dSTINGΔ* mutant cells ACC concentrated in the perinuclear region with minimal signal in the cell periphery (**Figure 8A**). The expression of *GFP-dSTING* on *dSTING* null background normalized ACC staining towards wild-type distribution. Calnexin staining was not affected by the mutation. Closer examination of the perinuclear region of fat body cells revealed that in *dSTINGΔ* mutants, ACC appeared disorganized and aggregated as compared to wild type cells and cells expressing *GFP-dSTING* (**Figure 8B**). The quantifications showed that 67% of nuclei had a perinuclear “aggregated” ACC phenotype (**Figure 8C**). On the other hand, FAS maintained its localization pattern in *dSTINGΔ* mutant cells (**Figure 9A**), but partially co-localized with ACC “aggregates” (**Figure 9B**), in agreement with immunoprecipitation results (**Figure 5C,D**).

**Figure 8.**
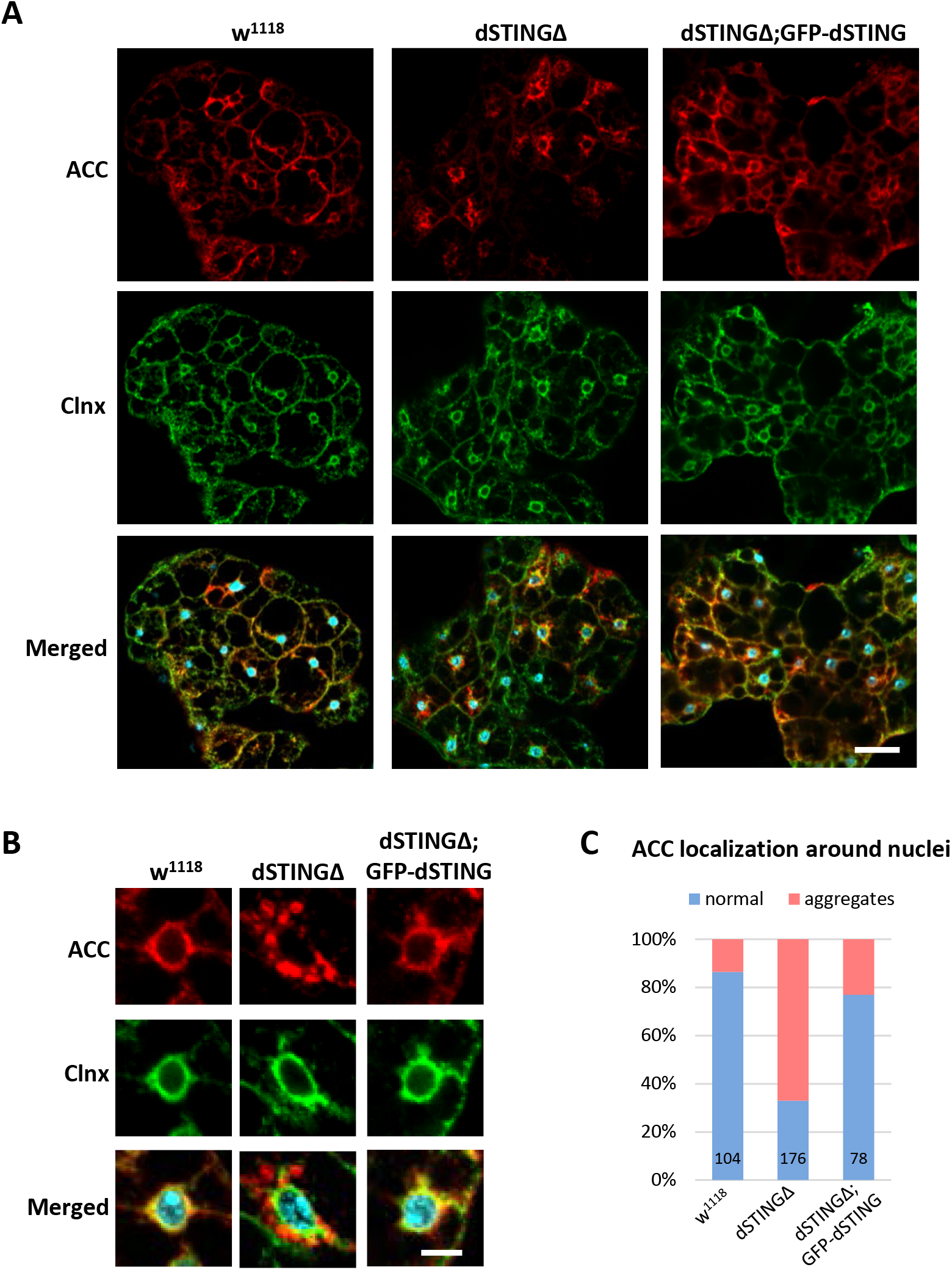
ACC localization is perturbed in *Drosophila STING* mutant fat body. **(A)** ACC has decreased cortical localization in *dSTINGΔ* mutant fat body as compared to control (*w*^*1118*^) and “rescue” (*dSTINGΔ;GFP-dSTING*) fly strains. Scale bar 20μm. **(B)** ACC localization in perinuclear region of fat body cells. Scale bar 5μm. **(C)** Quantification of perinuclear ACC localization pattern. Number of nuclei analyzed is shown for each genotype. Adult fat bodies were stained with ACC (red), Calnexin (Clnx, green) and DAPI (blue).

**Figure 9.**
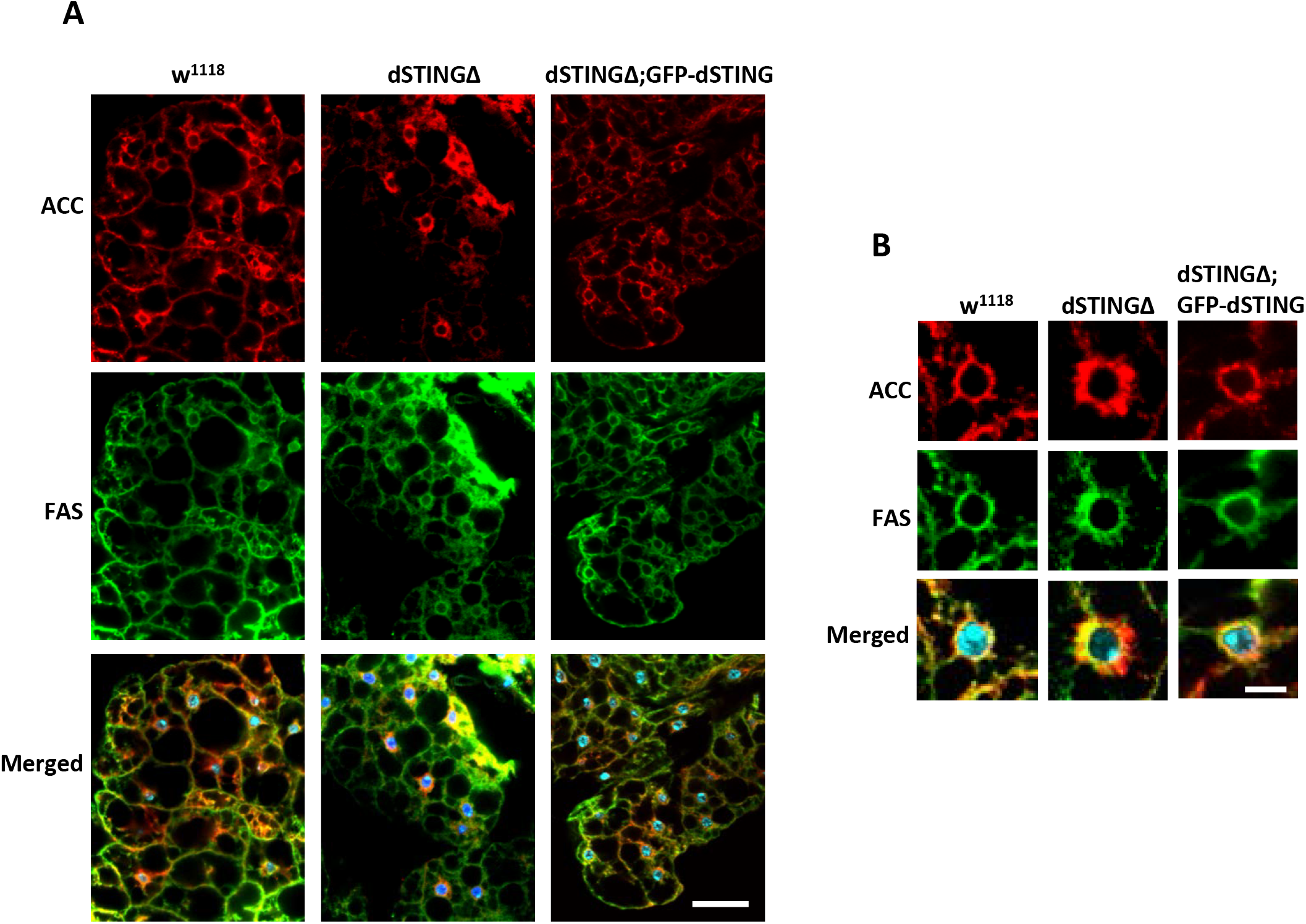
FAS localization is not affected in *dSTING*Δ mutant fat body. **(A)** FAS localization is not changed in *dSTINGΔ* mutant fat body as compared to control (*w*^*1118*^) and “rescue” (*dSTINGΔ;GFP-dSTING*) fly strains. Scale bar 20μm. **(B)** FAS and ACC localization in perinuclear region of fat body cells. Scale bar 5μm. Adult fat bodies were stained with ACC (red), FAS (green) and DAPI (blue).

Thus, we conclude that the presence of dSTING in fat body cells is required for proper ACC localization. In the absence of dSTING, ACC is no longer able to localize at the cell periphery and forms aggregated structures around fat body nucleus. FAS localization is mostly unchanged in *dSTINGΔ* mutants.

## DISCUSSION

Stimulator of interferon genes (STING) plays an important role in innate immunity of mammals, where activation of STING induces type I interferons (IFNs) production following the infection with intracellular pathogens (Ishikawa and Barber 2008) (Ishikawa *et al.* 2009) (Sun *et al.* 2009) (Tanaka and Chen 2012) (Zhong *et al.* 2008). However, recent studies showed that the core components of STING pathway evolved more than 600 million years ago, before the evolution of type I IFNs (Wu *et al.* 2014) (Margolis *et al.* 2017; Morehouse *et al.* 2020). This raises the question regarding the ancestral functions of STING. In this study we found that STING protein is involved in lipid metabolism in *Drosophila*. The deletion of *Drosophila STING (dSTING)* gene rendered flies sensitive to starvation and oxidative stress. These flies had reduced lipid storage and downregulated expression of lipid metabolism genes. We further showed that dSTING interacted with lipid synthesizing enzymes acetyl-CoA carboxylase (ACC) and fatty acid synthase (FAS) suggesting possible regulatory role in lipid biosynthesis. In fat body, main lipogenic organ in *Drosophila*, dSTING co-localized with both ACC and FAS in cortical region of ER. *dSTING* deletion resulted in disturbed ACC localization in fat body cells, and greatly reduced activity of FAS in *in vitro* assay.

Importantly, we also observed that ACC and FAS interacted with each other. Malonyl-CoA - the product of ACC - serves as a substrate for FAS reaction of fatty acid synthesis. Enzymes that are involved in sequential reactions often physically interact with each other and form larger multi-enzyme complexes, which facilitate substrate channeling and efficient regulation of pathway flux (Schmitt and An 2017), (Kastritis and Gavin 2018), (Sweetlove and Fernie 2018) (Zhang and Fernie 2020). There are several evidences of the existence of the multi-enzyme complex involved in fatty acid biosynthesis. ACC, ACL (ATP citrate lyase) and FAS physically associated in the microsomal fraction of rat liver (Gillevet and Dakshinamurti 1982). Moreover, in the recent work, lipogenic protein complex including ACC, FAS and four more enzymes was isolated from an oleaginous fungus *C. bainieri* (Shuib *et al.* 2018). It is possible that similar multi-enzyme complex exists in *Drosophila* and other metazoan species, and it would be of great interest to identify its other potential members.

How does STING exerts its effect on lipid synthesis? Recently, the evidence has emerged for the control of the *de novo* fatty acid synthesis by two small effector proteins - MIG12 and Spot14. MIG12 overexpression in livers of mice increased total fatty acid synthesis and hepatic triglyceride content (Kim *et al.* 2010). It has been shown that MIG12 protein binds to ACC and facilitates its polymerization thus enhancing the activity of ACC (Kim *et al.* 2010) (Park *et al.* 2013). For Spot14, both activation and inhibition of *de novo* lipogenesis have been reported, depending upon the tissue type and the cellular context (Rudolph *et al.* 2014), (Lafave *et al.* 2006), (Knobloch *et al.* 2013). Importantly, there is an evidence that all four proteins - ACC, FAS, MIG12 and Spot14 - exist as a part of a multimeric complex (Mckean 2016). It is plausible to suggest that *Drosophila* STING plays role similar to MIG12 and/or Spot14 in regulating fatty acid synthesis. We propose that dSTING might “anchor” ACC and FAS possibly together with other enzymes at the ER membrane. The resulting complex facilitates fatty acid synthesis by allowing for quicker transfer of malonyl-CoA product of ACC to the active site of FAS. In *dSTINGΔ* mutants, ACC loses its association with some regions of ER resulting in weakened interaction between ACC and FAS. We did observe less FAS immunoprecipitated with ACC in *dSTINGΔ* mutants compared to control flies, and the opposite effect was found in flies expressing GFP-tagged dSTING **(Figure 5C**).

It has been shown that *de novo* synthesis of fatty acids continuously contributes to the total fat body TAG storage in *Drosophila* (Heier and KÜhnlein 2018) (Wicker-Thomas *et al.* 2015) (Parvy *et al.* 2012) (Garrido *et al.* 2015). We hypothesize that reduced fatty acid synthesis due to the lowered FAS enzyme activity in *dSTINGΔ* deletion mutants might be responsible for the decreased TAG lipid storage and starvation sensitivity phenotypes. Sensitivity to oxidative stress might also be explained by reduced TAG level. Evidences exist that lipid droplets (consisting mainly of TAGs) provide protection against reactive oxygen species (Bailey *et al.* 2015) (Jarc *et al.* 2018) (Liu *et al.* 2015a). Furthermore, flies with ACC RNAi are found to be sensitive to oxidative stress (Katewa *et al.* 2012).

In addition to its direct role in ACC/FAS complex activity, STING might also affect phosphorylation status of ACC and/or FAS. Both proteins are known to be regulated by phosphorylation/dephosphorylation (Horton *et al.* 2002), (Boone *et al.* 2006), (Tong 2005), (Jin *et al.* 2010). In mammals, STING is an adaptor protein that transmits upstream signal by interacting with kinase TBK1 (TANK-binding kinase 1). When in complex with STING, TBK1 activates and phosphorylates IRF3 allowing its nuclear translocation and transcriptional response (Tanaka and Chen 2012) (Liu *et al.* 2015b) (Zhong *et al.* 2008). It is possible that in *Drosophila*, STING recruits a yet unidentified kinase that phosphorylates ACC and/or FAS thereby changing their enzymatic activity.

*Drosophila* STING itself could also be regulated by the lipid- synthesizing complex. STING palmitoylation was recently identified as a posttranslational modification necessary for STING signaling in mice (Mukai *et al.* 2016) (Hansen *et al.* 2019) (Hansen *et al.* 2018). In this way, palmitic acid synthesized by FAS might participate in the regulation of dSTING possibly providing a feedback loop.

The product of ACC - malonyl-CoA - is a key regulator of energy metabolism (Saggerson 2008). During lipogenic conditions, ACC is active and produces malonyl-CoA, which provides the carbon source for the synthesis of fatty acids by FAS. In *dSTING* knockout, FAS activity is decreased and malonyl-CoA is not utilized and builds up in the cells. Malonyl-CoA is also a potent inhibitor of carnitine palmitoyltransferase CPT1, the enzyme that controls the rate of fatty acid entry into the mitochondria, and hence is a key determinant of the rate of fatty acid oxidation (Mcgarry and Brown 1997). Thus, high level of malonyl-CoA results in decreased fatty acid utilization for energy. This might explain the down-regulation of lipid catabolism genes that we observed in *dSTINGΔ* mutants (**Figure 4E**). Reduced fatty acid oxidation in turn shifts cells to the increased reliance on glucose as a source of energy. Consistent with this notion, we observed increased glucose level in fed *dSTINGΔ* mutant flies (**Figure 2D**), as well as increased levels of phosphoenolpyruvate (PEP) (**Supplementary Figure 7)**. PEP is produced during glycolysis and its level was shown to correlate with the level of glucose (Moreno-Felici *et al.* 2019). Reliance on glucose for energy also has a consequence of reduced incorporation of glucose into trehalose and glycogen for storage, and therefore, lower levels of these storage metabolites, which we observed (**Figure 2B,C**). Based on our findings we propose a model summarized in **Figure 10**, which suggests a direct involvement of dSTING in the regulation of lipid metabolism.

**Figure 10.**
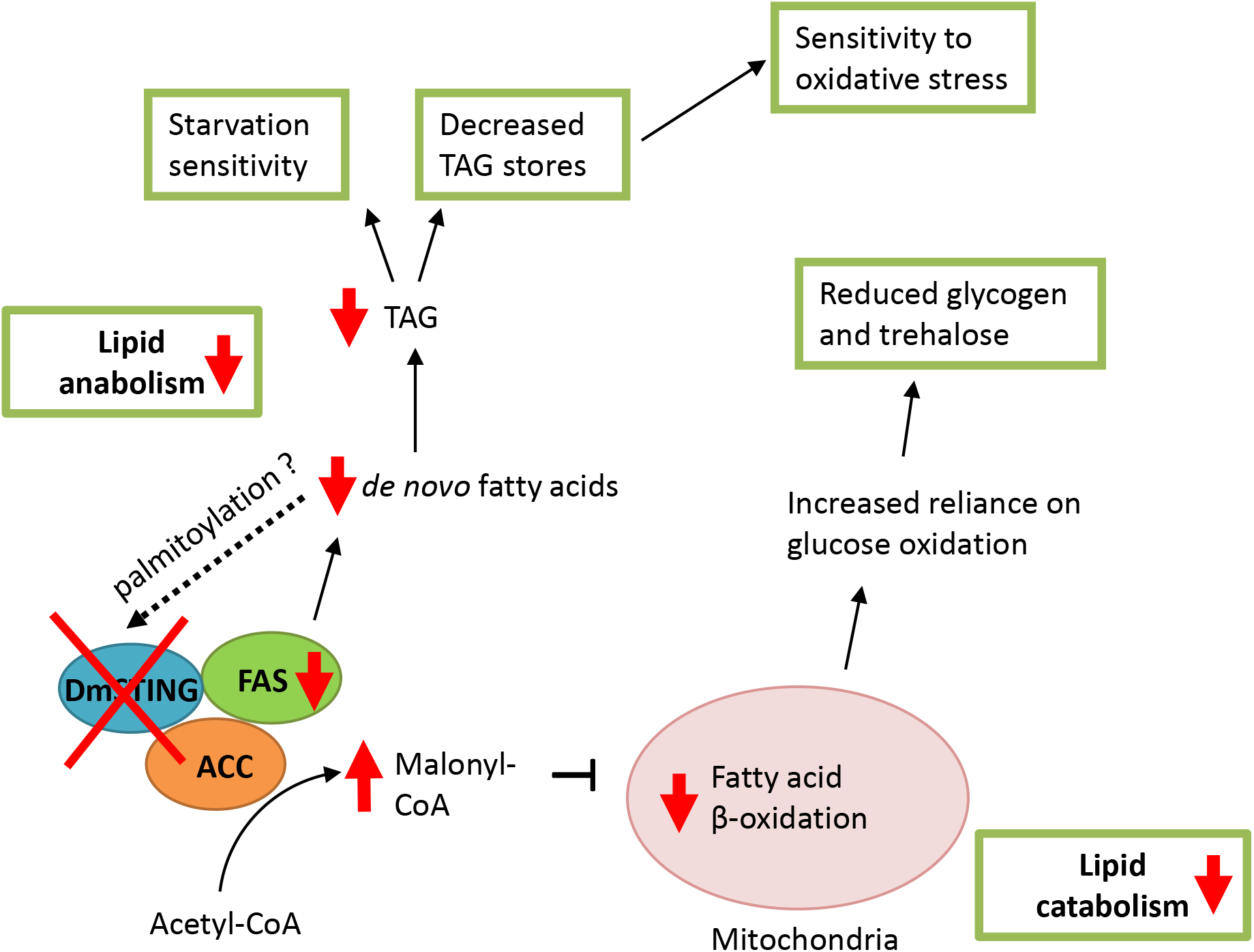
Model of *dSTING* deletion effect on *Drosophila* metabolism. See text for details.

Recent studies show that in mammals, the STING pathway is involved in metabolic regulation under obesity conditions. The expression level and activity of STING were upregulated in livers of mice with HFD (high fat diet)-induced obesity (Bai *et al.* 2017). STING expression was increased in livers from NAFLD (nonalcoholic fatty liver disease) patients compared to control group (Luo *et al.* 2018). In NASH (nonalcoholic steatohepatitis) mouse livers, STING mRNA level was also elevated (Xiong *et al.* 2019). Importantly, STING deficiency ameliorated metabolic phenotypes and decreased lipid accumulation, inflammation and apoptosis in fatty liver hepatocytes (Iracheta-Vellve *et al.* 2016) (Petrasek *et al.* 2013) (Qiao *et al.* 2018).

Despite the accumulating evidences, exact mechanism of STING functions in metabolism is not completely understood. The prevailing hypothesis is that obesity leads to mitochondrial stress and subsequent mtDNA release into the cytoplasm, which activates cGAS-STING pathway (Bai *et al.* 2017) (Bai and Liu 2019) (Yu *et al.* 2019). The resulting chronic sterile inflammation is responsible for the development of NAFLD, insulin resistance and type 2 diabetes. In this case, the effect of STING on metabolism is indirect and is mediated by inflammation effectors. The data presented in the current study strongly suggest that in *Drosophila*, STING protein is directly involved in lipid metabolism by interacting with enzymes involved in lipid biosynthesis. This raises the question if the observed interaction is unique for *Drosophila* or it is also the case for mammals. Future work is needed to elucidate the evolutionary aspect of STING role in metabolism. Understanding the relationships between STING and lipid metabolism may provide insights into the mechanisms of obesity-induced metabolism dysregulation and thereby suggest novel therapeutic strategies for metabolic diseases.

## MATERIALS AND METHODS

### Fly stocks

Deletion mutations of *dSTING* gene (*dSTINGΔ*) were created by imprecise excision of *P* element-based transposon *P{EPgy2}Sting*^*EY06491*^(FBti0039337). This transposon is mapped 353 bp upstream of the *dSTING* start codon. To initiate excision, males *y1,w*^*67c23*^*; P{w*^+*mC*^*, y*^+*mDint2*^=*EPgy2} STING*^*EY0649*^(Bloomington stock 16729) were crossed to females of the “jump” stock *y*^*1*^*w*^*1118*^;*CyO, PBac*{*w*+^mC^*=Delta 2-3. Exel*}*2*/*amos*^*Tft*^, bearing Δ2-3 transposase on a second chromosome, marked by *Curly*. F1 *Curly* males *y*^*1*^*w*^*1118*^; *P{w*^+*mC*^*, y*^+*mDint2*^=*EPgy2} STING*^*EY0649*^/*CyO, PBac*{*w+mC*=*Delta 2-3. Exel*}*2* were collected and crossed to *w*^*1118*^; *If*/*CyO* females. The resulting F2 progeny was screened for white-eyed flies. White-eyed flies were crossed individually to *w*^*1118*^; *If*/*CyO* to set up stocks *dSTINGΔ*/*CyO* and then *w*^*1118*^; *dSTINGΔ/dSTINGΔ* homozygotes. The genomic DNA of these mutants was isolated. Mutations were confirmed by sequence determination following the PCR amplification with *dSTINGΔ* primer: 5’-CTCAGAATTCTCATTTATTCTGGCC -3’. RT-PCR analysis of *dSTING* expression confirmed that obtained deletions are *dSTING* null mutations.

For rescue experiments, *pCasper*-based vector containing UAS sequence followed by native *dSTING* promoter (437 bp upstream *dSTING* start codon) and GFP-tagged *dSTING* cDNA (clone #LP14056, DGRC, Bloomington) was injected into *w*^*1118*^ *Drosophila* embryos (Model System Injections, Raleigh, NC). Fly stocks *w*^*1118*^*; dSTINGΔ/dSTINGΔ; GFP-dSTING-WT/GFP-dSTING-WT* were set up. The expression of tagged proteins was verified by immunoblot analysis with anti-GFP antibody.

Fly stock *y*^*1*^*v*^*1*^*;P{TRiP.HMJ23183}attP40/CyO* (NIG-Fly, National Institute of Genetics, Japan) was used for RNAi experiments.

For overexpression or RNAi in fat body, *cg-GAL4* driver was used (Bloomington stock 7011). For ubiquitous overexpression in mass-spec experiment, *tub-GAL4* driver was used (Bloomington stock 5138). Fly stock *y^1^v^1^;P{TRiP.HMJ23183}attP40/CyO* (NIG-Fly, National Institute of Genetics, Japan) was used for RNAi experiments together with *cg-GAL4* driver.

For genetic interaction with *foxo* experiments, fly stock with *foxo*^*Δ94*^ deletion was used (Bloomington stock 42220).

Flies were grown and maintained on food consisting of the following ingredients: 1 part of Nutri-Fly^®^ GF (Genesee Scientific, cat. 66-115) and 3 parts of Jazz-mix (Fisher Scientific, cat. AS153). All crosses were carried out at 25°C. *w*^*1118*^ fly stock was used as a wild-type control.

### Starvation and oxidative stress

For life span analysis, newly enclosed flies (females or males) were transferred to fresh food every 2 days, and dead flies were counted.

For starvation stress assay, 5-days old adult flies (females or males) were transferred from normal food to the vials containing Whatman filter paper soaked with PBS (15-20 flies per vial). Fresh PBS was added every 24 hours to prevent drying. Dead flies were counted every 12 hours.

For oxidative stress assay, 5-days old adult flies (females or males) were transferred from normal food to the vials containing normal food supplemented with 5% hydrogen peroxide (15-20 flies per vial). Dead flies were counted every 12 hours.

For starvation and oxidative stress resistance experiments on larvae, 2^nd^ instar larvae (~53 hours) were transferred to media containing 1.2% agarose (starvation) or 1.2% agarose with 10% sucrose and 10mM paraquat (oxidative stress). Survived larvae were counted every 12 hours.

### Axenic flies

To obtain axenic flies, 0-12 hour embryos were collected, dechorionated for 5 min in 50% Clorox, washed 2x with autoclaved water and transferred to sterile food. The axenity of flies was confirmed by PCR from flies homogenate using primers to 16s rDNA gene (8FE, 5’- AGAGTTTGATCMTGGCTCAG-3’ and 1492R, 5’- GGMTACCTTGTTACGACTT-3’).

### Triglycerides and glycogen quantifications

Eight 5-days old males (with heads removed) were collected, frozen in liquid nitrogen and stored at −80°C. Flies were ground in 200μl of PBST buffer (PBS with 0.01% Triton X-100) and heated at 70°C for 10 minutes (Tennessen *et al.* 2014).

For TAG measurement, 6μl of homogenate were mixed with 25μl of PBS and 30μl of TAG reagent (Pointe Scientific, Cat. T7531) or Free Glycerol Reagent (MilliporeSigma, Cat. F6428). Triglyceride standard solution (from Pointe Scientific, Cat. T7531 kit) and glycerol standard solution (MilliporeSigma, Cat. G7793) were used as standards. Reactions were incubated for 30 minutes at 37°C, centrifuged 6000g for 2 minutes and supernatants were transferred to 96-well plate, after which absorbance was read at 540nm. The TAG concentration in each sample was determined by subtracting the values of free glycerol in the corresponding sample. Total protein level in the samples was determined using Bio-Rad Protein Assay Dye Reagent Concentrate (Bio-Rad, Cat. 5000006).

For glycogen measurement, homogenate was centrifuged 5 minutes at 10000g. 6μl of supernatant were mixed with 24μl of PBS and 100μl of glucose reagent (MilliporeSigma, Cat. GAGO20) with or without the addition of amyloglucosidase (MilliporeSigma, Cat. A1602, 0.25U per reaction) and transferred to 96-well plate. Glycogen solution (Fisher Scientific, Cat. BP676-5) and glucose solution (MilliporeSigma, Cat. 49161) were used as standards. Reactions were incubated 60 minutes at 37°C, after which 100μl of sulfuric acid were added to stop the reaction, and the absorbance was read at 540nm. Glycogen concentration in each sample was determined by subtracting the values of free glucose in corresponding sample. Total protein level in the samples was determined using Bio-Rad Protein Assay Dye Reagent Concentrate (Bio-Rad, Cat. 5000006).

### Hemolymph sugars quantification

Fifty 5-days old males were anesthetized with CO_2_ and pricked with a needle in the thorax. 0.2 ml PCR tubes with caps removed were inserted inside 1.5 ml tube. Pricked flies were placed into a spin column (Zymo Research, Cat. N. C1005-50) with plastic ring and filling removed (leaving only bottom glass wool layer). Spin columns were inserted into a 1.5 ml tube with PCR tube, centrifuged 5 min at 2500 g at 4°C, shaken to dislodge flies and centrifuged one more minute. 0.5 μl of collected hemolymph were mixed with 4.5 μl of PBS, heated at 70°C for 5 min, centrifuged at 6000g 15 sec and placed on ice. To measure glucose level, 2 μl samples (in duplicates) were mixed with 100μl Infinity glucose reagent (Thermo Scientific, Cat. N. TR15421) in a 96-well plate, and after 5 min incubation at 37°C the absorbance was read at 340 nm. To measure trehalose level, 1μl of trehalase (MilliporeSigma, Cat. No T8778) was added to the wells with measured glucose (see above). Plate was incubated at 37°C overnight, the absorbance was read at 340 nm, and glucose readings were subtracted from obtained values. Total protein level in the samples was determined using Bio-Rad Protein Assay Dye Reagent Concentrate (#5000006, Bio-Rad).

### Capillary feeder assay (CAFE) assay

Capillary feeder (CAFE) assay was adopted from (Diegelmann *et al.* 2017). Plastic bottles with carton caps and small holes on the bottom to allow for air circulation were used. Five openings were made in a carton cap to fit the pipette tips of 2-20 μl volume. Five glass capillaries (Drummond Scientific Company, Cat. No. 2-000-001) were filled with 5 μl of 20% sucrose solution in water and inserted into pipet tips on the cap. Ten 4-days old males were placed in each bottle, and all bottles were placed into a plastic box containing wet paper towel to provide humidity. Control bottles that contained no flies were set up to account for liquid evaporation. After 24 hours and 48 hours the amount of food consumed in each bottle was measured as follows: Food consumption (μl/fly) = (Food uptake (μl) - Evaporative loss (μl)) / total number of flies in the vial.

### Smurf gut permeability assay

5-days old flies were transferred from normal food to food containing 2.5% (wt/vol) Blue dye no. 1 (MilliporeSigma, Cat. No 3844-45-9). Flies were kept on dyed food for 12 h. A fly was counted as a Smurf if dye coloration could be observed outside of the digestive tract.

### Nile Red staining

Adult fat body and guts were dissected in PBS, fixed in 4% paraformaldehyde for 20 minutes, washed twice with PBS and mounted in fresh Nile Red solution with DAPI (0.5 mg/ml Nile Red stock solution diluted 1000x with PBS supplemented with 30% glycerol). Images were collected using Olympus Fluoview FV3000. Quantification of lipid droplets area was performed using cellSens Dimension Desktop (Olympus). Minimum eight samples per genotype were analyzed.

### Microarray analysis

Drosophila Gene 1.0 ST CEL files were normalized to produce gene-level expression values using the implementation of the Robust Multiarray Average (RMA) (Irizarry 2003) in the *affy* package (version 1.48.0) (Gautier *et al.* 2004) included in the Bioconductor software suite (version 3.2) (Gentleman *et al.* 2004) and an Entrez Gene-specific probeset mapping (20.0.0) from the Molecular and Behavioral Neuroscience Institute (Brainarray) at the University of Michigan (Dai 2005). Array quality was assessed by computing Relative Log Expression (RLE) and Normalized Unscaled Standard Error (NUSE) using the *affyPLM* package (version 1.46.0).

Principal Component Analysis (PCA) was performed using the *prcomp* R function with expression values that had been normalized across all samples to a mean of zero and a standard deviation of one. Differential expression was assessed using the moderated (empirical Bayesian) *t* test implemented in the *limma* package (version 3.26.9) (i.e., creating simple linear models with *lmFit*, followed by empirical Bayesian adjustment with *eBayes*). Correction for multiple hypothesis testing was accomplished using the Benjamini-Hochberg false discovery rate (FDR) (Benjamini *et al.* 2001). Human homologs of fly genes were identified using HomoloGene (version 68). All microarray analyses were performed using the R environment for statistical computing (version 3.2.0).

Gene Ontology (GO) analysis was conducted using the DAVID Functional Annotation Tool (https://david.ncifcrf.gov/).

Gene Set Enrichment Analysis (GSEA) (version 2.2.1) (Subramanian *et al.* 2005) was used to identify biological terms, pathways and processes that are coordinately up- or down-regulated within each pairwise comparison. The Entrez Gene identifiers of the human homologs of the genes interrogated by the array were ranked according to the *t* statistics computed for each effect in the two-factor model and for each pairwise comparison. Any fly genes with multiple human homologs (or *vice versa*) were removed prior to ranking, so that the ranked list represents only those human genes that match exactly one fly gene. Each ranked list was then used to perform pre-ranked GSEA analyses (default parameters with random seed 1234) using the Entrez Gene versions of the Hallmark, Biocarta, KEGG, Reactome, Gene Ontology (GO), and transcription factor and microRNA motif gene sets obtained from the Molecular Signatures Database (MSigDB), version 6.0 (Subramanian *et al.* 2007).

### RT-qPCR

RNA was isolated from eight 5 days old males using ZR Tissue and Insect RNA MicroPrep™ (Zymo Research, #R2030). DNA was removed using TURBO™ DNase (Invitrogen, #AM2238) following manufacturer’s recommendations. cDNA was generated from 1ug of total RNA using ProtoScript^®^ II First Strand cDNA Synthesis Kit (New England Biolabs, E6560). RT-qPCR analysis was performed in Luna Universal qPCR Master Mix (New England Biolabs, #M3003) using a Roche LightCycler480 (Roche). Primers used were: *bmm* (PP222440), *rpl32* (PD41810), *InR* (PP37075), *Thor/4E-BP* (PD43730) (Hu *et al.* 2013). Two qPCR technical replicates were conducted for three biological replicates. Relative expression was normalized to *rpl32* reference gene using ΔΔCt comparative method.

### Nuclear and Cytoplasmic Proteins Extraction

Nuclear and cytoplasmic fractions were extracted from twenty 5 days-old males following the existing protocol (Piccolo *et al.* 2015) and subjected to Western blotting. Flies were either fed or starved for 24 hours on PBS. Antibodies used were: Gapdh1 (Sigma-Aldrich, #G9545, 1:2000), β-tubulin (1:1000, DSHB, E7-c), Foxo (1:1000, Abcam, ab195977).

### Mass spectrometry

Fat body from six third instar larvae ubiquitously overexpressing GFP-dSTING (genotype *w*^*1118*^*;+/+;tub-GAL4/GFP-dSTING*) or control larvae (genotype *w*^*1118*^) were ground in 200μl of IP buffer (25mM HEPES, pH 7.6, 0.1 mM EDTA, 12 mM MgCl_2_, 100mM NaCl, 1% NP-40) and extracted for 30 min at RT. Recombinant GFP protein was added to the control lysate. Samples were centrifuged 10,000g for 5 min and supernatant was precleared with 20μl of protein G sepharose beads (Amersham Biosciences, cat. 17-0618-01) for 2 hours at 4°C. Precleared lysate was incubated with 4 μg of antibodies against GFP tag (DSHB, 4C9) overnight at 4°C. Beads were washed four times with IP buffer, and immunoprecipitation reactions were separated by SDS-PAGE and most prominent individual gel bands corresponding to ~250kDa and ~30kDa were excised. Mass spectrometry detection was performed at The Proteomics Resource Center at The Rockefeller University. Proteins were reduced with DTT, alkylated with iodoacetamide and trypsinized. Extracted peptides were analyzed by nanoLC-MS/MS (Dionex 3000 coupled to Q-Exactive+, Thermo Scientific), separated by reversed phase using an analytical gradient increasing from 1% B/ 99% A to 40% B/ 60% A in 27 minutes (A: 0.1% formic acid, B: 80% acetonitrile/0.1% formic acid). Identified peptides were filtered using 1% False Discovery Rate (FDR) and Percolator (KÄll *et al.* 2007). Proteins were sorted out according to estimated abundance. The area is calculated based on the most abundant peptides for the respective protein (Silva *et al.* 2006). Proteins not detected or present in low amounts are assigned an area zero. Data were extracted and queried against Uniprot *Drosophila* using Proteome Discoverer and Mascot.

### Immunoprecipitation

Fifteen abdomens of 5 days old males were ground in 300μl of IP buffer (10mM Tris pH 7.4, 1mM EDTA, 1mm EGTA, 2mM MgCl_2_, 2mM MnCl_2_, 1 x Halt Protease and Phosphatase Inhibitor Cocktail (Thermo Scientific, cat. 78446), supplemented with 100mM NaCl, 0.02% Triton X-100 for ACC IP and 150mM NaCl, 0.1% Triton X-100 for FAS IP). After extraction for 30 min at RT samples were centrifuged 600g 3 min and supernatants were precleared with 15μl of protein A agarose beads (Goldbio, cat. P-400-5) for 2 hours at RT. After discarding the beads, supernatant was divided in half and incubated with either antibodies or corresponding normal IgG overnight at 4°C. Antibodies used were: rabbit anti-ACC (Cell Signaling, #3676), guinea pig anti-FAS (generously provided by A.Teleman (Moraru *et al.* 2018)), rabbit IgG (Sino Biological, cat. CR1), guinea pig IgG (Sino Biological, cat. CR4). Beads were washed three times with IP buffer, and bound proteins were analyzed by SDS-PAGE and Western blotting.

### Acetyl-CoA carboxylase activity assay

Assay was conducted using Acetyl-CoA Carboxylase assay kit (#MBS8303295, MyBioSource). Eight males were collected, frozen in liquid nitrogen and stored at −80°C. Flies were ground in 250μl of assay buffer after which another 250μl of assay buffer were added (total lysate volume 500μl). Lysates were centrifuged 8,000g for 10 min at 4°C, and 300μl of supernatant were transferred to a new tube. To set up the reaction, 10μl of supernatant (or assay buffer for control reactions) were mixed with 90μl of substrate and incubated 30 min at 37°C after which the reactions were centrifuged 10,000g for min at 4°C. 5μl of supernatant, water (for blank reaction) or standards (phosphate) were added to 100μl of dye working reagent in a 96-well plate, and the absorbance at 635nm was recorder after 5 min of incubation. Total protein level in the samples was determined using Bio-Rad Protein Assay Dye Reagent Concentrate (#5000006, Bio-Rad). One unit of Acetyl-CoA Carboxylase activity is defined as the enzyme generates 1 nmol of PO_4_^3-^per hour.

### Fatty acid synthase activity assay

Assay was conducted essentially as described in (Moraru *et al.* 2018). Eight males were collected, frozen in liquid nitrogen and stored at −80°C no more than one day. Flies were ground in 150μl of homogenization buffer (10mM potassium phosphate buffer pH 7.4, 1mM EDTA, 1mM DTT) and 300μl of cold saturated ammonium sulfate solution (4.1M in water, pH 7) were added to the lysate. After incubation on ice for 20 min, samples were centrifuged at 20,000g for 10 min at 4°C and supernatant was carefully removed. Pellet was resuspended in 200μl of homogenization buffer, centrifuged 10,000g 10 min, and 150μl of supernatant were transferred to a new tube. To set up the reaction, 20μl of sample were added to 160 μl of 0.2mM NADPH (#9000743, Cayman Chemical) in 25mM Tris pH 8.0 and incubated 10 min at 25°C in a 96-well plate. 20μl of water (for control reaction) or a mix of 10μl of 0.66mM acetyl CoA (#16160, Cayman Chemical) and 10μl of 2mM malonyl-CoA (#16455, Cayman Chemical) were added to the reaction, and absorbance at 340 nm was recorded every 5 min for 60 min at 25°C using Synergy 2 multi-mode microplate reader (BioTec). Absorbance for control reaction were subtracted for each time point. Total protein level in the samples was determined using Bio-Rad Protein Assay Dye Reagent Concentrate (#5000006, Bio-Rad).

### Polar metabolite profiling

For polar metabolite profiling experiment, twenty 5 days old adult flies (males) were collected in Eppendorf tube, weighted, frozen in liquid nitrogen and stored at −80°C. For metabolite extraction, flies were transferred to 2ml tubes with 1.4 mm ceramic beads (Fisher Scientific, cat. 15-340-153), 800μl of extraction buffer were added (80% methanol (A454, Fisher Scientific), 20% H_2_0 (W7SK, Fisher Scientific), standards) and flies were processed on BeadBlaster™ 24 Microtube Homogenizer (Benchmark Scientific) at 6 m/s for 30 s. Tubes were incubated on rotator for 1h at 4°C and centrifuged 20,000g 15 min at 4°C. 700μl of supernatant were transferred to new Ependorf tube and dried in a vacuum centrifuge.

Metabolomics analysis was performed at The Proteomics Resource Center at The Rockefeller University. Polar metabolites were separated on a ZIC-pHILIC 150 × 2.1 mm (5-μm particle size) column (EMD Millipore) connected to a Thermo Vanquish ultrahigh-pressure liquid chromatography (UPLC) system and a Q Exactive benchtop orbitrap mass spectrometer equipped with a heated electrospray ionization (HESI) probe. Dried polar samples were resuspended in 60 μl of 50% acetonitrile, vortexed for 10 s, centrifuged for 15 min at 20,000g at 4°C and 5 μl of the supernatant was injected onto the LC/MS system in a randomized sequence. Mobile phase A consisted of 20 mM ammonium carbonate with 0.1% (vol/vol) ammonium hydroxide (adjusted to pH 9.3) and mobile phase B was acetonitrile. Chromatographic separation was achieved using the following gradient (flow rate set at 0.15 ml min−1): gradient from 90% to 40% B (0–22 min), held at 40% B (22–24 min), returned to 90% B (24–24.1 min), equilibrating at 90% B (24.1–30 min). The mass spectrometer was operated in polarity switching mode for both full MS and data-driven aquisition (DDA) scans. The full MS scan was acquired with 70,000 resolution, 1 × 106 automatic gain control (AGC) target, 80-ms max injection time and a scan ranges of 110-755 m/z (neg), 805-855 m/z (neg) and 155-860 m/z (pos). The data-dependent MS/MS scans were acquired at a resolution of 17,500, 1 × 105 AGC target, 50-ms max injection time, 1.6-Da isolation width, stepwise normalized collision energy (NCE) of 20, 30 and 40 units, 8-s dynamic exclusion and a loop count of 2, scan range of 110-860 m/z.

Relative quantitation of polar metabolites was performed using Skyline Daily56 (v.20.1.1.158) with the maximum mass and retention time tolerance were set to 2 ppm and 12 s, respectively, referencing an in-house library of chemical standards. Metabolite levels were normalized to the total protein amount for each condition.

### Membrane and cytoplasmic Proteins Extraction

Membrane fractionation was performed following the protocol from (Abas and Luschnig 2010) with modifications. Flies were either fed or starved for 24 hours. Thirteen abdomens (guts and testes removed) from 5 days old males of flies expressing GFP-dSTING (*w*^*1118*^*;dSTINGΔ/dSTINGΔ;GFP-dSTING/GFP-dSTING*) were ground in 100μl of EB (30mM Tris pH 7.5, 25% sucrose, 5% glycerol, 5mM EDTA, 5mM EGTA, 5mM KCl, 1mM DTT, aprotinin, leupeptin, PMSF), spun down at 600g for 3 min to remove debris. Supernatant after centrifugation represents total protein fraction. Supernatant was diluted twice with 100 μl H_2_O and centrifuged at 21,000g for 2 hours at 4°C. Resulting supernatant represents cytoplasmic fraction. Pellet was resuspended in 30μl of EB supplemented with 0.5% Triton X-100, resulting in membrane fraction sample. Proteins were subjected to SDS-PAGE and Western blotting. Total protein fraction was used for assessing the levels of ACC and FAS. Cytoplasmic and membrane fractions were used to analyze GFP-dSTING localization. Antibodies used were: ACC (1:1000, C83B10, Cell Signaling, #3676), FAS (1:2000, generously provided by A.Teleman (Moraru *et al.* 2018)), Gapdh1 (1:2000, Sigma-Aldrich, #G9545), ATPβ (1:1000, Abcam, cat. ab14730), GFP (1:1000, Santa Cruz Biotechnology, B2, cat. sc-9996).

### Immunostaining

Adult fat body and guts were dissected in PBS, fixed in 4% paraformaldehyde for 20 minutes, washed with PBST (PBS supplemented with 0.1% Triton X-100) and blocked with PBST supplemented with 10% goat serum for 1 hour at RT. Tissue were stained with primary antibodies in PBST + 10% goat serum overnight at 4°C, washed three times with PBST, and incubated with secondary antibodies in PBST + 10% goat serum for 2 hours at RT. Antibodies used were: ACC (1:200, C83B10, Cell Signaling, #3676), Calnexin (1:30, DSHB, Cnx99A, 6-2-1-s), FAS (1:150, generously provided by A.Teleman (Moraru *et al.* 2018)), GFP (Proteintech, cat. 50430-2-AP). After three washes with PBST, tissues were stained with DAPI, washed with PBS and mounted in Fluoromount-G (SouthernBiotech, cat. 0100-01). Images were collected using Olympus Fluoview FV3000.

### Lysotracker staining

Fat body of fed or starved early third instar larvae was dissected in PBS, incubated for 5 minutes in 100μM LysoTracker Red DND-99 (Invitrogen, L7528) in PBS and mounted in PBS supplemented with 30% glycerol. Images were taken using Olympus BX61 motorized upright microscope fitted with a BX-DSU disc scan unit.

## ACKNOWLEDGMENTS

We would like to thank Adam Gower from the Boston University Microarray and Sequencing Resource Core Facility. All research materials and data from our studies will be freely available to other investigators. This work was supported by a grant from NIH to IC (GM121449).

## Supplementary figures

**Supplementary Figure 1.**
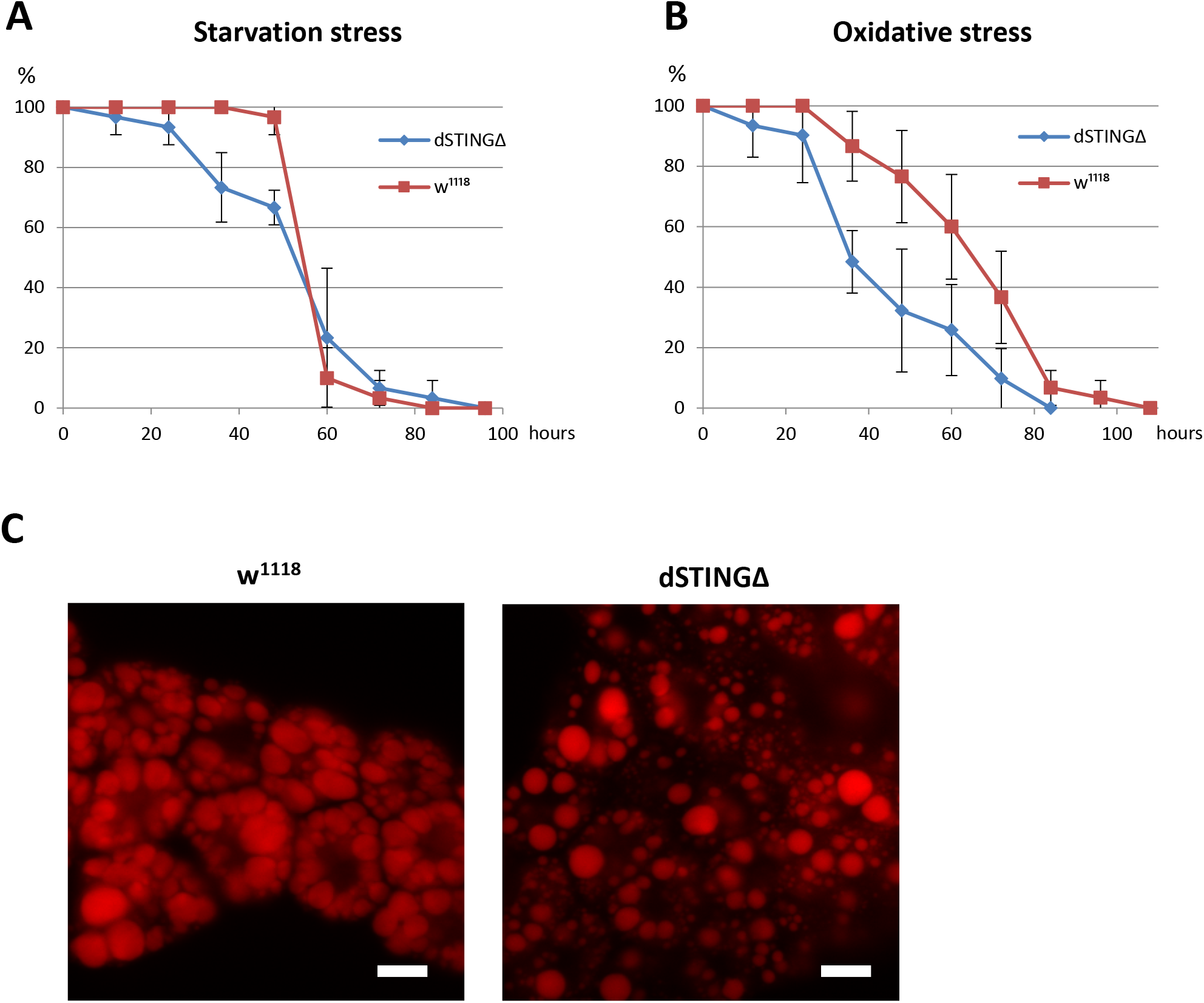
*Drosophila STING* mutant larvae are susceptible to starvation and oxidative stress and have decreased lipid storage. **(A)** Starvation resistance. 2^nd^ instar larvae (~53 hours) were transferred to media containing 1.2% agarose. Survived larvae were counted every 12 hours. n = 30. **(B)** Oxidative stress resistance. 2^nd^ instar larvae (~53 hours) were transferred to media containing 1.2% agarose, 10% sucrose and 10mM paraquat. Survived larvae were counted every 12 hours. n = 30. **(C)** Lipid staining. Fat body of 24 hour-starved early third instance larvae were stained with Nile Red. Scale bar 10 μm.

**Supplementary Figure 2.**
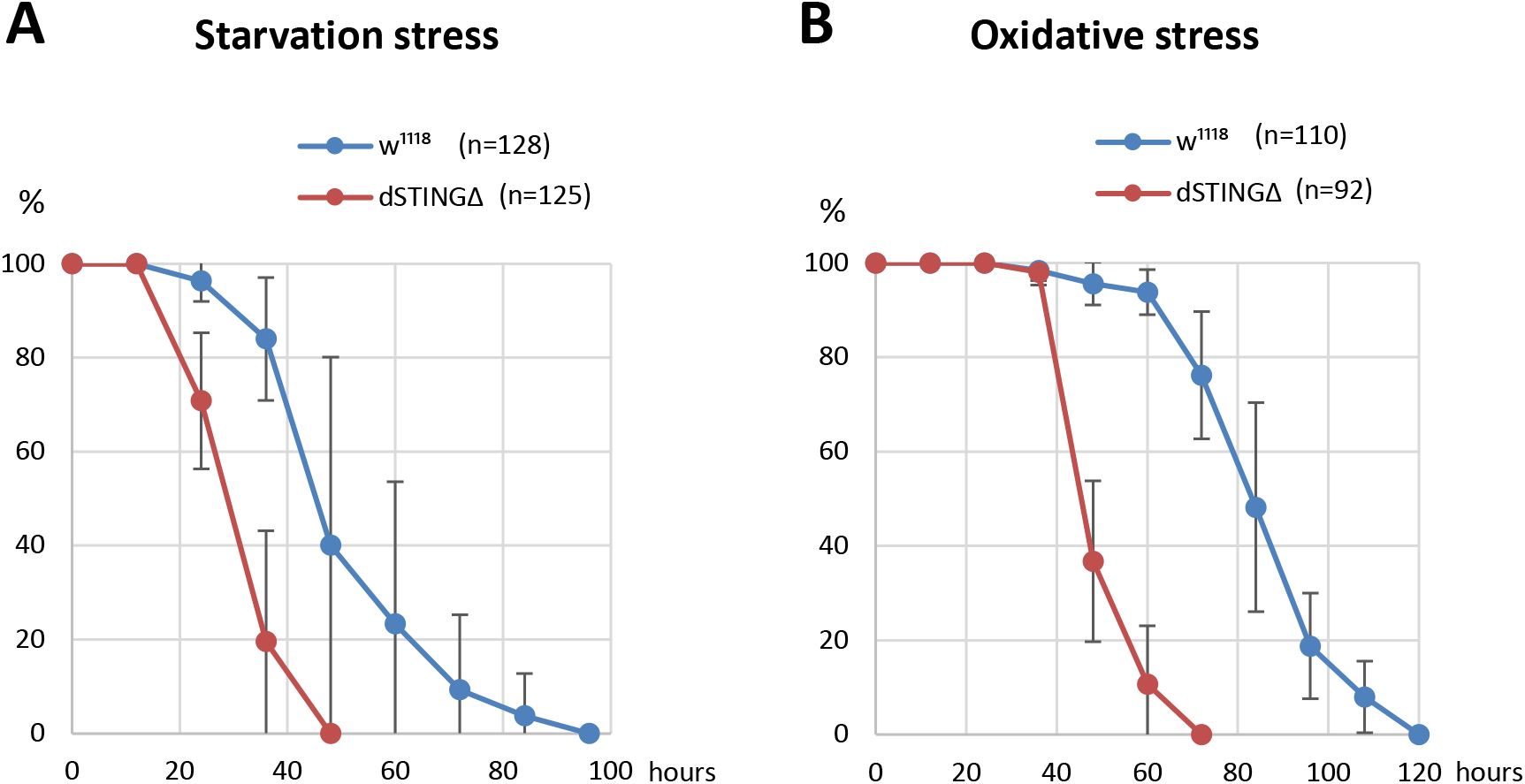
*Drosophila STING* mutants are susceptible to starvation and oxidative stress under axenic condition. **(A)** Starvation stress resistance of axenic males. Flies were kept on PBS only and survived flies were counted every 12 hours. **(B)** Oxidative stress resistance of axenic males. Flies were kept on food supplemented with 5% hydrogen peroxide and survived flies were counted every 12 hours. Percentages of survived flies at each time point are shown. The number of flies analyzed is shown in chart legend for each genotype.

**Supplementary Figure 3.**
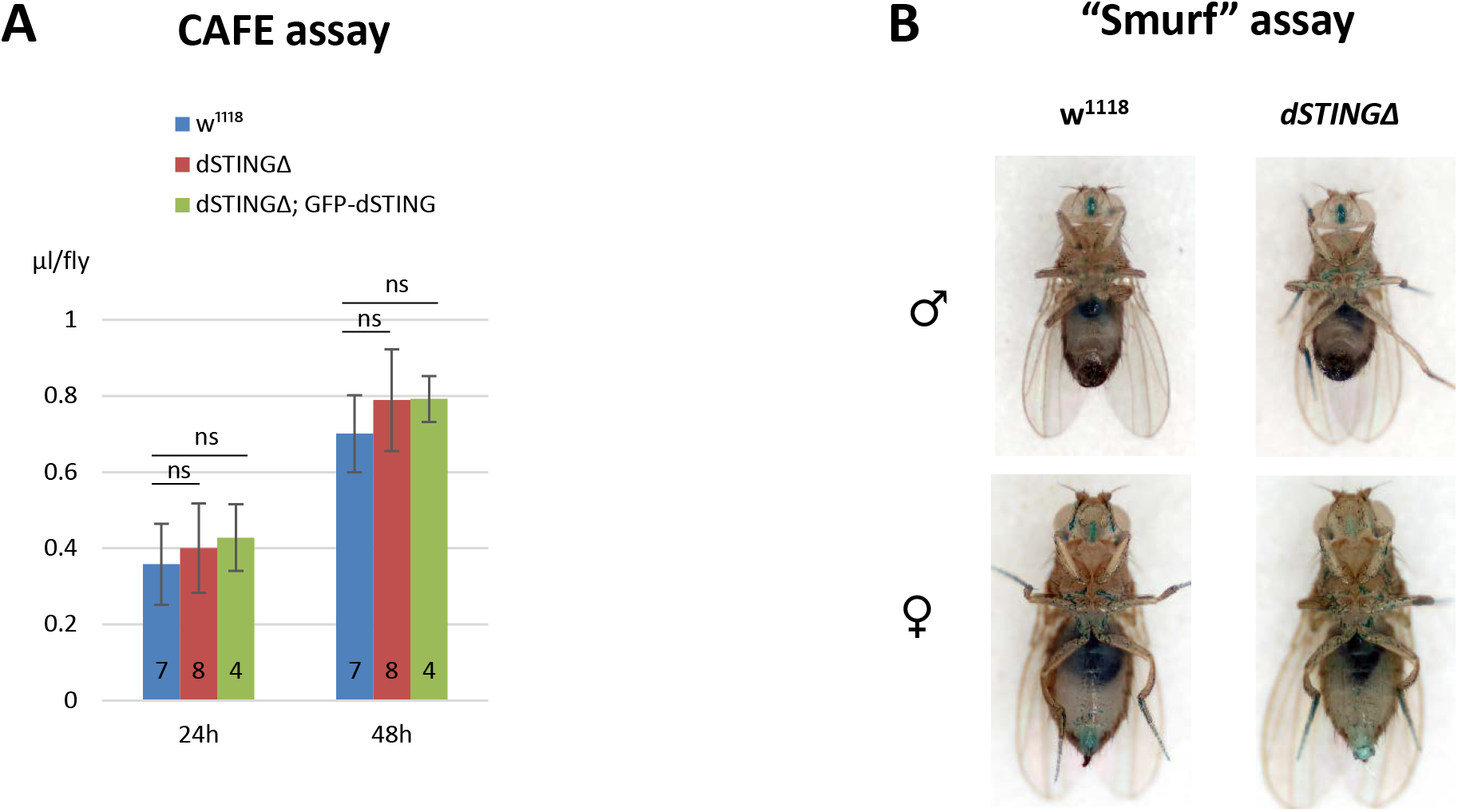
Food ingestion is not compromised in *Drosophila STING* mutant flies. **(A)** Capillary feeder (CAFE) assay. The amount of food consumed by each fly is shown during time period indicated. The number of experiments for each genotype is indicated. Student’s t-test, ns indicates statistically non significant. **(B)**“Smurf” assay. Flies were kept on food containing blue dye for 12h. Unabsorbable blue dye did not penetrate the intestinal wall in both wild type and *dSTINGΔ* mutants.

**Supplementary Figure 4.**
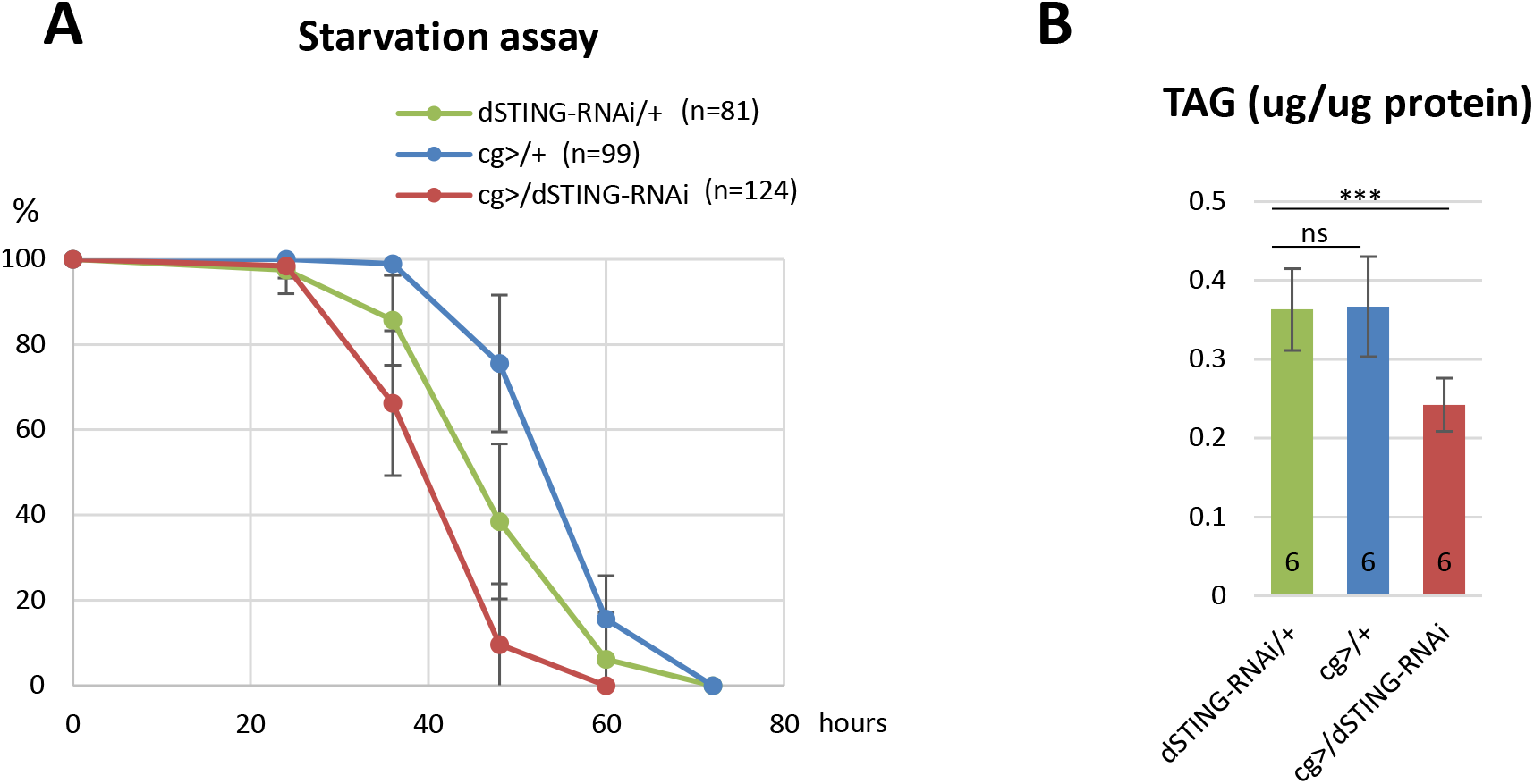
Fat body specific RNAi of *dSTING* results in increased sensitivity to starvation and decreased TAG level. **(A)** Starvation stress resistance of males. Flies were kept on PBS only and survived flies were counted every 12 hours. Percentages of survived flies at each time point are shown. The number of flies analyzed is shown in chart legend for each genotype. **(B)** Total body TAG level. The number of experiments for each genotype is indicated. Student’s t-test, ***p<0.001, ns indicates statistically non significant. *dSTING* RNAi was induced specifically in fat body with *cg-GAL4* driver. Following strains were used in the experiment: *dSTING-RNAi*/+ [RNAi control], *cg>/+* [driver control], *cg>/dSTING-RNAi* [RNAi of *dSTING* in fat body].

**Supplementary Figure 5.**
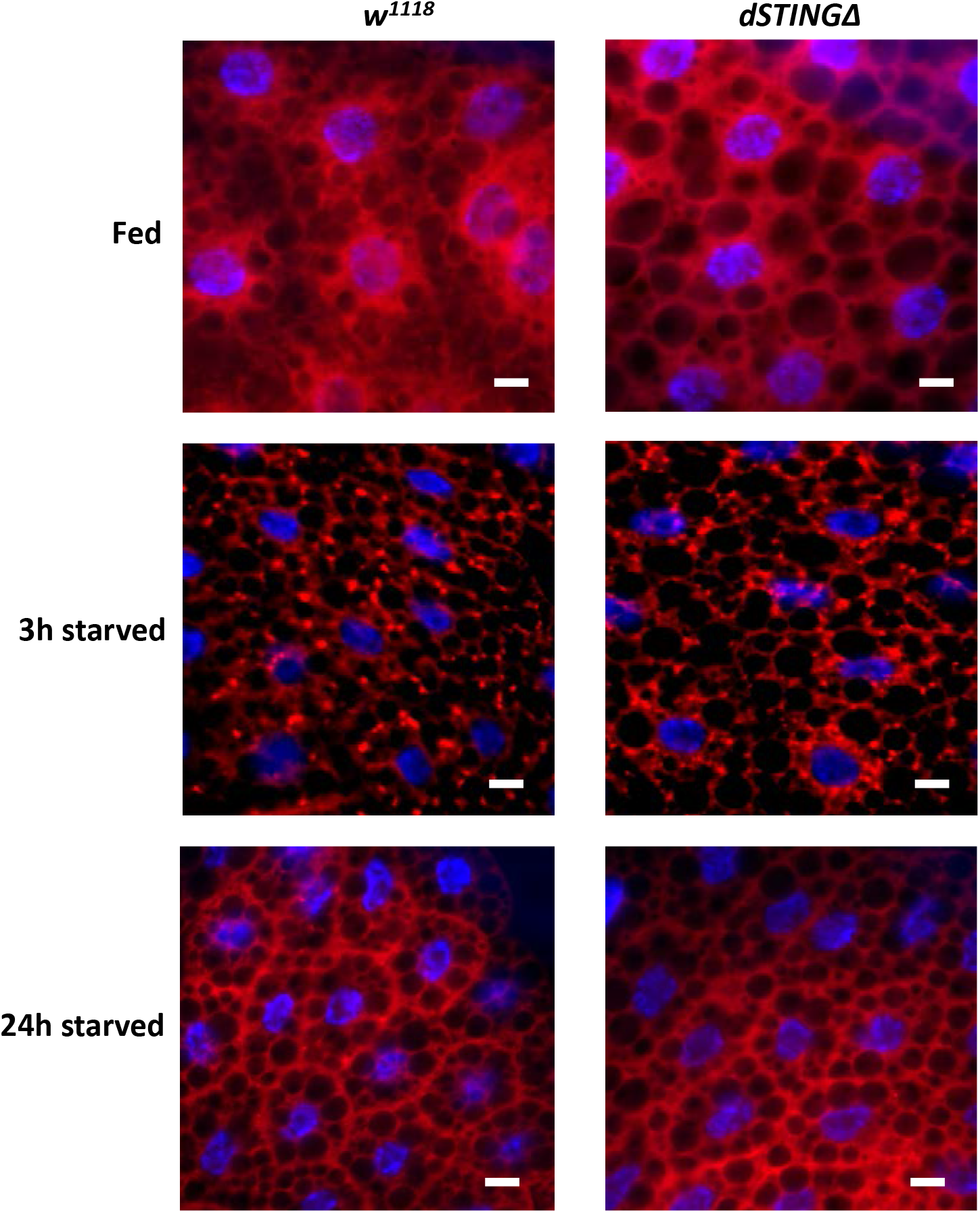
Starvation-induced autophagy is not perturbed in *Drosophila STING* mutants. Early 3^rd^ instar larvae were starved for 3 hours or 24 hours, and their fat bodies were stained with Lysotracker (red). Nuclei were stained with DAPI (blue). After 3 hours of starvation autophagy is induced in both control (*w1118*) and mutants (*dSTINGΔ*). After 24 hours of starvation the autophagy response is cleared in both control and *dSTINGΔ* mutants. Scale bar 10μm.

**Supplementary Figure 6.**
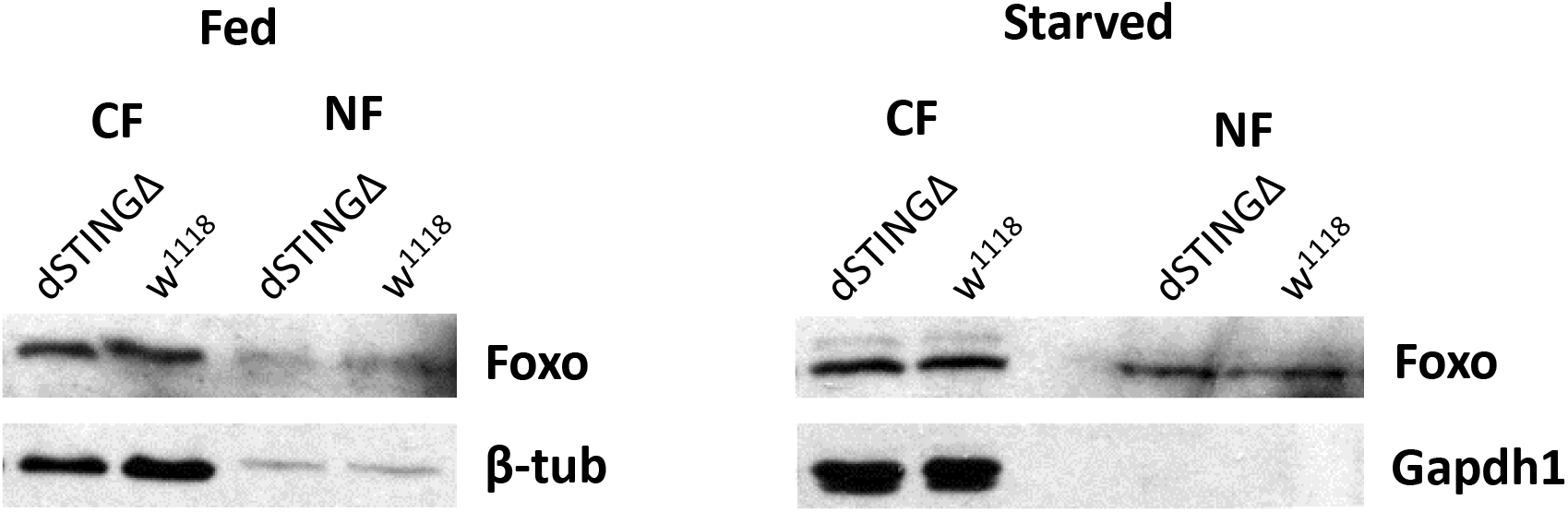
Nuclear FoxO is not increased in *dSTINGΔ* mutants. Nuclear (NF) and cytoplasmic (CF) fractions were extracted from 5 days-old males and subjected to Western blotting. Flies were either fed or starved for 24 hours.

**Supplementary Figure 7.**
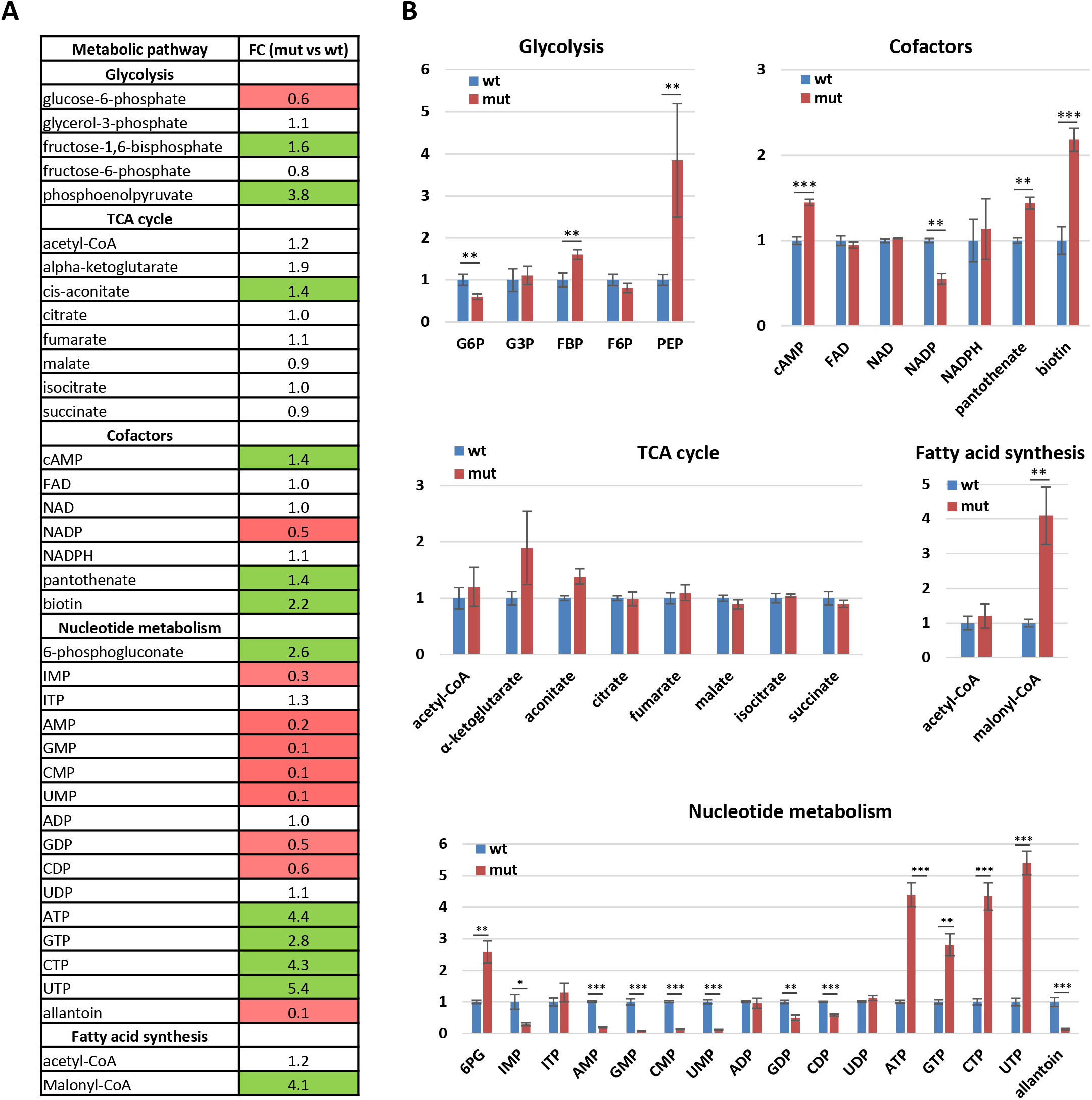
Metabolomics analysis of *Drosophila STING* mutants. **(A)** List of metabolites identified. Left column shows the names of metabolites and metabolic pathway they relate to. Right column shows fold change (FC) values (mut vs wt, where “mut” corresponds to *dSTINGΔ* mutant “wt” to *w*^*1118*^, respectively). Colored cells represent statistically significant difference, where red indicates decreased level, green indicates increased level. **(B)** Charts show changes in metabolite levels in *dSTINGΔ* mutants (mut, red) relative to control *w*^*1118*^ (wt, blue). Student’s t-test, ***p<0.001, **p<0.01, *p<0.1.

**Supplementary Figure 8.**
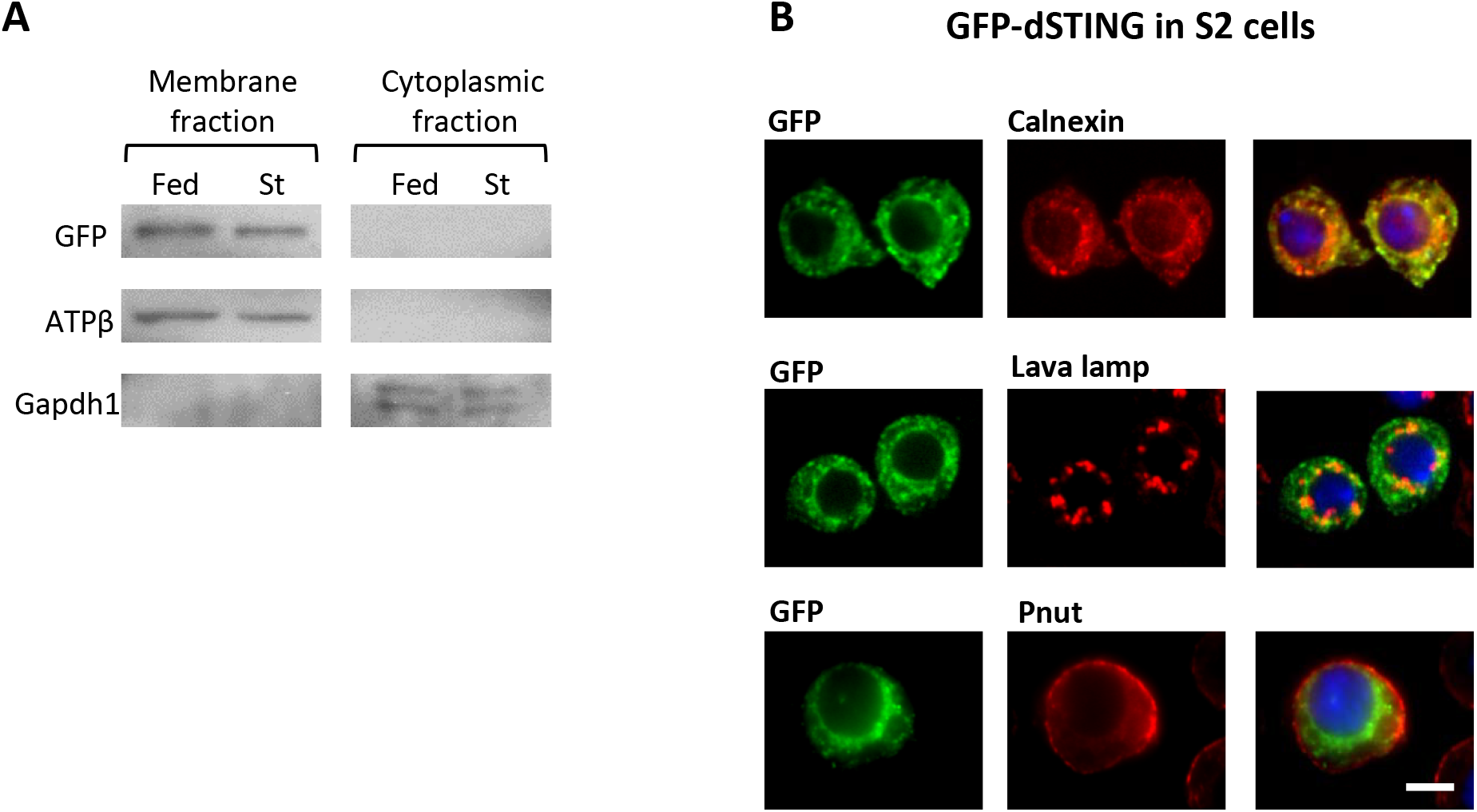
*Drosophila* STING localizes at the endoplasmic reticulum membrane. **(A)** GFP-dSTING is associated with membrane fraction. Membrane and cytoplasmic fractions were extracted from abdomens of five days old males (of genotype *dSTINGΔ;GFP-dSTING*) and subjected to Western blotting. Flies were either fed or starved for 24 hours. ATPβ and Gapdh1 were used as markers for membrane and cytoplasmic fractions, respectively. **(B)** GFP-dSTING is localized at endoplasmic reticulum (ER). *Drosophila S2* tissue culture cells expressing GFP-dSTING were immunostained with following antibodies: GFP, Calnexin (ER marker), Lava lamp (Golgi marker), Pnut (cellular membrane marker). Nuclei were stained with DAPI. Scale bar 5 μm.

## Notes

### Competing Interest Statement

The authors have declared no competing interest.

